# Bacterial bioconvection confers context-dependent growth benefits and is robust under varying metabolic and genetic conditions

**DOI:** 10.1101/2021.04.20.440374

**Authors:** Daniel Shoup, Tristan Ursell

## Abstract

Microbial communities often respond to environmental cues by adopting collective behaviors—like biofilms or swarming—that benefit the population. Bioconvection is a distinct and robust collective behavior wherein microbes locally gather into dense groups and subsequently plume downward through fluid environments, driving flow and mixing on scales thousands of times larger than an individual cell. Though bioconvection was observed more than 100 years ago, effects of differing physical and chemical inputs, as well as its potential selective advantages to different species of microbes, remain largely unexplored. In the canonical microbial bioconvector *Bacillus subtilis*, density inversions that drive this flow are setup by vertically oriented oxygen gradients that originate from an air-liquid interface. In this work, we develop *Escherichia coli* as a complementary model organism for the study of bioconvection. We show that for *E. coli* and *B. subtilis*, bioconvection confers a context-dependent growth benefit with clear genetic correlates to motility and chemotaxis. We found that fluid depth, cell concentration, and carbon availability have complimentary effects on the emergence and timing of bioconvective patterns, and whereas oxygen gradients are required for *B. subtilis* bioconvection, we found that *E. coli* deficient in aerotaxis (Δ*aer*) or energy-taxis (Δ*tsr*) still bioconvect, as do cultures that lack an air-liquid interface. Thus, in two distantly related microbes, bioconvection confers context-dependent growth benefits, and *E. coli* bioconvection is robustly elicited by multiple types of chemotaxis. These results greatly expand the set of physical and metabolic conditions in which this striking collective behavior can be expected and demonstrate its potential to be a generic force for behavioral selection across ecological contexts.

## Introduction

In complex natural environments, as well as relatively simple fluids, populations of microbes both shape and respond to the chemical and physical landscapes that impose selective pressures (1–11). For instance, microbial communities in aquatic ecosystems often reside in relatively still fluid environments at scales where both flow and diffusion are relevant transport processes (11–15). In response to a persistent and vertically aligned gradient, cells can ascend the fluid column en masse to produce mass-density inversions that drive large-scale flow and advection, aptly termed ‘bioconvection’ (16). The resulting distribution of cell density is spatially complex and temporally dynamic, but trends in the temporal and spatial evolution of such patterned flow are reproducible products of physical and chemical factors (1, 17–22). This phenomenon was first examined in freshwater protozoa nearly 60 years ago (23) and had been observed decades earlier when initially well-mixed cultures spontaneously segregated into visible regions of high and low cell density (24, 25).

The basic mechanism is well demonstrated by the canonical microbial bioconvector *Bacillus subtilis.* In liquid cultures, aerobic *B. subtilis* sense and ascend oxygen gradients (26) that emanate from air-liquid interfaces (so-called ‘aerotaxis’) (18, 27). Cells collect in dense groups near air-liquid interfaces (28, 29) such that the spatial distribution of cell density stores gravitational potential energy. Under the influence of gravity, these groups—and the corresponding mass-density inversion—inevitably succumb to a Rayleigh-Taylor instability in which dense plumes of bacteria drift downward, driving convective flow (17, 29, 30), distinct from propulsive flow generated by the cells themselves (31). The ecological relevance of bioconvection is amplified by the fact that bioconvection is not uniquely a product of oxygen gradients; multiple environmental cues, such as light and gravity also produce global, vertically aligned stimuli that foster bioconvective behavior (21, 32). The same basic mechanisms drive bioconvection in the protozoa *Tetrahymena* (23) and algae in the genus *Chlamydomonas* (20, 33).

Bioconvection can also be characterized by the combination of physical parameters that elicit it (34, 35). First, a non-equilibrium process (e.g., differential heating or chemotaxis) creates biased motion and subsequent mass-density variations that store gravitational potential energy in the fluid. Second, gravity acts on those density variations to produce flow that transports material primarily along the axis that aligns with gravity. Third, diffusion of the relevant actors or molecules reduces density gradients that store gravitational potential energy. Fluid viscosity modifies both the effective diffusion coefficient (e.g., through the Stokes-Einstein relation) and the rate at which convection dissipates potential energy into the fluid as heat. Lastly, increasing fluid depth increases the length-scale over which density variations and diffusion compete, with deeper fluids strongly favoring convection. These parameters assemble to form the dimensionless *Rayleigh number R* = Δ*ρgL*^3^/*Dμ*, where Δ*ρ* is the scale of mass-density variations, *g* is the acceleration gravity, *L* is the fluid depth, *D* is the diffusion coefficient, and *μ* is the fluid viscosity (36). Assuming that the viscosity does not depend on the presence or behaviors of the cells at relevant concentrations (37–39), the parameters *g*, *μ*, and *L* have no biological dependencies. This leaves complexities of cellular motion, behavior, metabolism, and the evolution of attendant cellular and chemical concentration fields as inputs to Δ*ρ* and *D*. The Rayleigh number captures the onset of convection by comparing two characteristic time scales. The first is the time scale on which diffusion equilibrates the density variations along the fluid column, *τ*_1_ = *L*^2^/*D*. The second is a time scale inversely related to the rate at which the gravity-driven instability develops, *τ*_2_ = *μ*/Δ*ρgL*. If the Rayleigh number (*R* = *τ*_1_/*τ*_2_) is less than one, diffusion dominates and convection is inhibited. If the Rayleigh number is near one, the fluid convects in stable laminar patterns (as in Rayleigh-Bénard convection (40)). If the Rayleigh number is much larger than one, turbulent convection emerges (41, 42).

Modeling dynamics in such systems is extraordinarily complex, as it involves both ‘typical’ fluid dynamics that now must incorporate time-and-space varying fluid density and the spatiotemporal evolution of both agent and chemical fields that diffuse, interact, and advect with the motion of the fluid. Nevertheless, a number of previous studies have successfully reproduced key attributes of this phenomena (21, 27, 43–48) across multiple species (49), as well as expanding models to include a thermal gradient (50). In addition to the balance of time scales, the qualitative understanding discussed above suggests that if one considers an aqueous, Newtonian fluid (viscosity independent of shear (51)) in a terrestrial environment, necessary conditions for bioconvection include (i) exogenous flow must be much slower than flow generated by convection, (ii) populations of cells must move in response to chemical (or other) gradients; and (iii) nutrient concentrations must be able to support a motile population of sufficient concentration. These conditions are sufficiently generic to indicate that bioconvection is likely a ubiquitous, albeit hard to observe, phenomenon across many species, with the potential to produce ecologically important flows that move chemical and biological material from the scale of bacteria up to the scale of macroscopic aquatic organisms (13).

However, these conditions do not address other salient questions: Which types of chemotaxis can elicit bioconvection? Does bioconvection require an air-liquid interface and the attendant oxygen gradient? How do metabolic inputs affect bioconvection? As they migrate, cells consume metabolites en masse with the potential to alter the chemical landscape that guides subsequent chemotactic migration (52–55). This suggests that organisms that respond to the dynamic structure of the chemical landscape (as opposed to the static structure of light gradients or gravity) might exhibit bioconvective patterns that are sensitive to the presence of specific nutrients. *B. subtilis*, however, is not an ideal species in which to test this hypothesis because the order in which it consumes nutrients via metabolism is not directly correlated with its chemotactic preference for those nutrients (56) and because it primarily responds to the oxygen gradient generated by a stationary air-liquid interface (18, 22). On the other hand, *Escherichia coli* exhibit chemotactic sensitivity to multiple metabolites that aligns with their preferred order of consumption (56, 57). This direct correlation between metabolite consumption and chemotaxis suggested that *E. coli* could be used to interrogate how changes in nutrient availability, or the absence of a persistent oxygen gradient, influence microbial bioconvection.

The goals of this work are (i) to assess whether bioconvection confers a selective advantage, (ii) to characterize (within in a limited range of parameter space) the effects of fluid depth and initial cell concentration on bioconvective patterns and timing, (iii) to assess the importance of different kinds of chemotaxis in eliciting bioconvection, and (iv) to interrogate nutrient and boundary conditions that facilitate and/or affect bioconvection. To address these questions, we developed *E. coli* as a versatile and robust model system for the study of microbial bioconvection, and we compared its behavior to that of the canonical microbial bioconvector, *B. subtilis*. First, we show that in *E. coli* and *B. subtilis*, bioconvective patterns require populations that are motile and chemotactic, albeit with differences between species, whereas run-only and tumble-only knock-out mutants of each species are bioconvection deficient. With those same knock-outs, we show that all three genotypes (wild-type, tumble-only, run-only) have essentially identical growth curves in shaken media, but when allowed to grow in still media, wild-type cells that bioconvect display a depth-dependent growth enhancement. We show that the presence of a simple sugar causes significant effects on the appearance and duration of bioconvective patterns and that aerotaxis-deficient (Δ*aer*) or energy-taxis-deficient (Δ*tsr*) *E. coli* can bioconvect, albeit with differences in patterns, indicating that direct oxygen sensing is not required for bacterial bioconvection. Finally, we show that *E. coli* produce bioconvective patterns in the absence of an air-liquid interface and the accompanying oxygen gradient. Together, our results demonstrate that *E. coli* is a robust model organism for studying bioconvection, with a wide range of physical and chemical conditions producing this ecologically important collective behavior. This work also suggests that in the context of bioconvection, the benefit of chemotaxis expands beyond whether an individual cell responds to its local chemical environment to include whether a population can respond en masse to elicit a collective behavior that benefits the population as a whole.

## Results

### Bioconvection confers context-dependent growth benefits in two distantly related species

The density inversions and large-scale alterations of chemical landscapes that drive bioconvection require movement of and consumption by large groups of cells, and hence bioconvection is a distinctly population-scale behavior. However, *B. subtilis* and *E. coli* respond differently to chemotactic signals in their environment; notably, *B. subtilis* exhibit strong attraction to oxygen (aerotaxis) (53) and therefore have been a model organism for the study of aerotaxis-mediated bioconvection (17, 18). *E. coli*, on the other hand, can grow anaerobically (58) and exhibit chemotaxis toward a diverse array of sugars, amino acids, and small molecules in their environment (26, 56, 57, 59–61). We wondered whether these chemotactic differences between species translate into observable differences in bioconvective group behaviors. We began by imaging the dynamic density patterns produced by bioconvection in these two canonical species and then determined whether the combination of motility and chemotaxis conferred similar, if any, growth benefits under conditions conducive to bioconvection.

We imaged bioconvective patterns using transmitted white-light microscopy. In this modality, an increase in the local density of bacteria scatters more light from the imaging path, producing darker pixel intensities, and conversely, where relatively few bacteria reside, the image appears lighter, thus producing intensity variations across the image that monotonically correlate with bacterial density (18). Spontaneous bacterial-density fluctuations initiated bioconvection and produced striking patterns as shown in Fig 1 (see also corresponding SI movies). These patterns and their evolution through time were a 2D projection of the full three- dimensional bacterial density field in a cylindrical sample. We started by visually comparing the patterns of *B. subtilis* and *E. coli*; for each species, we imaged wild-type (motile and chemotactic), run-only (motile but not chemotactic), and tumble-only (non-motile and not chemotactic) genotypes (see Methods). Wild-type *B. subtilis* formed laminar convection cells that maintained a roughly circular shape and whose diameter decreased over the course of 200 minutes (see SI video: Fig_1_BS_wt). In contrast, under identical conditions, wild-type *E. coli* cultures evolved through a range of visually distinct, turbulent convective patterns (see SI video: Fig_1_EC_wt), conceptually similar to pattern-transitions seen in other bioconvecting species (20, 62). For both species, wild-type genotypes produced intricate, dynamic, and high-contrast patterns that were absent in the respective run-only and tumble-only mutant strains. After hours of growth, run- only *B. subtilis* exhibited large-scale non-convective flow, potentially due to collective swimming (see SI video: Fig_1_BS_run-only); run-only *E. coli* displayed a slowly evolving sedimentation pattern (see SI video: Fig_1_EC_run-only). A combination of patterned cell sedimentation (63) and subtle dynamic patterning was observed in the tumble-only mutant strains of *E. coli* and *B. subtilis*, but neither exhibited convection patterns nor large-scale mixing (see SI videos: Fig_1_EC_tumble-only, Fig_1_BS_tumble-only).

**Figure 1.**
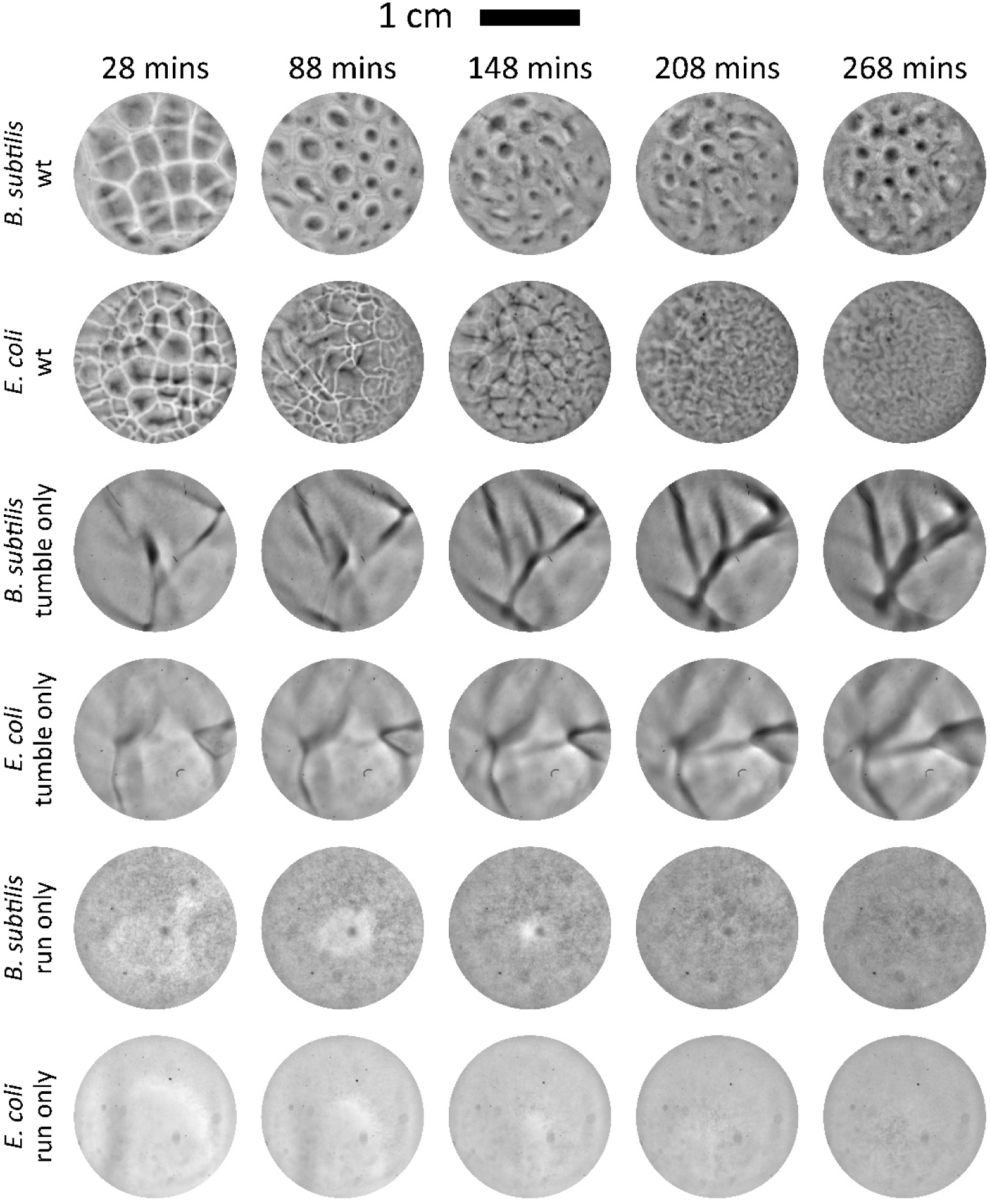
Bioconvective patterning and dynamics depend on chemotaxis and vary between *B. subtilis* and *E. coli*. To determine how bioconvection varies between *E. coli* and *B. subtilis*, cultures of wild-type, tumble-only, and run-only strains of each species were imaged under bioconvective conditions in LB media. Each strain was grown in shaking batch culture to OD 0.5 and placed in 12-well plates at a fluid depth of 2.5 mm. Top-down images were taken at the times indicated, and image intensity was approximately proportional to projected cell density. In each row, image contrast is shown saturated to 0.001% of the stack histogram, and intensities across each row are comparable. At early times, both wild-type *E. coli* and *B. subtilis* formed relatively large convection cells. *B. subtilis* formed consistent patterns for multiple hours, while *E. coli* evolved through multiple visually distinct patterns. Tumble-only strains of both species produced relatively static sedimentation patterns, while run-only strains of both species remained diffuse. See SI for corresponding movies.

Next, we wanted to know whether these genotypically dependent patterns of bioconvection corresponded to differences in the growth dynamics of their respective populations. We characterized the effects of bioconvection on *B. subtilis* and *E. coli* by comparing bulk growth rates of wild-type and knock-out mutants in both still (i.e., unshaken) and externally mixed (i.e., shaken) liquid cultures. In both species, our previous imaging (Fig 1) indicated that run-only and tumble-only mutant cultures were devoid of bulk flow driven by bioconvection. Thus, we hypothesized that any growth-rate enhancement due to bioconvection would be apparent between genotypes in a still environment, whereas no significant enhancement would be observed between genotypes in an externally mixed environment.

Starting with *B. subtilis,* we measured the bulk concentration of cells (OD) over time for individual cultures of wild-type, run-only, and tumble-only strains. In this series of experiments, cultures were grown in standard rich media (see Methods) within the cylindrical wells of 12-well plates at a fluid depth of 2.5 mm. First, we measured the average cell density (OD) of the entire culture as a function of time, in triplicate, for all genotypes in shaking culture conditions. Over 30 hours, the growth curves for each genotype in the shaken condition were statistically indistinguishable, having quantitatively similar growth rate, saturation time, and maximum density (Fig 2A). Then, we repeated this experiment in still cultures that allowed populations to adopt whatever bulk flows occurred naturally due to bacterial activity and/or bioconvection; all other conditions were identical, and controls indicated insignificant thermal flow (see Methods). Over the course of 35 hours, all three genotypes showed significant deficits in growth rate and maximum density as compared to shaken cultures, consistent with the fact that well-mixed cultures have higher levels of dissolved oxygen and therefore increased aerobic respiration (64). However, there *were* significant differences in growth rate, transient dynamics, and maximum density between the genotypes in a still physical context. Consistent with our hypothesis, wild-type cells that were both motile and chemotactic had the highest growth rate and attained the highest density, run-only cells had initial growth rates similar to wild-type but ultimately did not achieve the same maximum densities, and tumble-only cells had significant deficits in growth rate and maximum density as compared to wild-type and run-only genotypes (Fig 2B).

**Figure 2.**
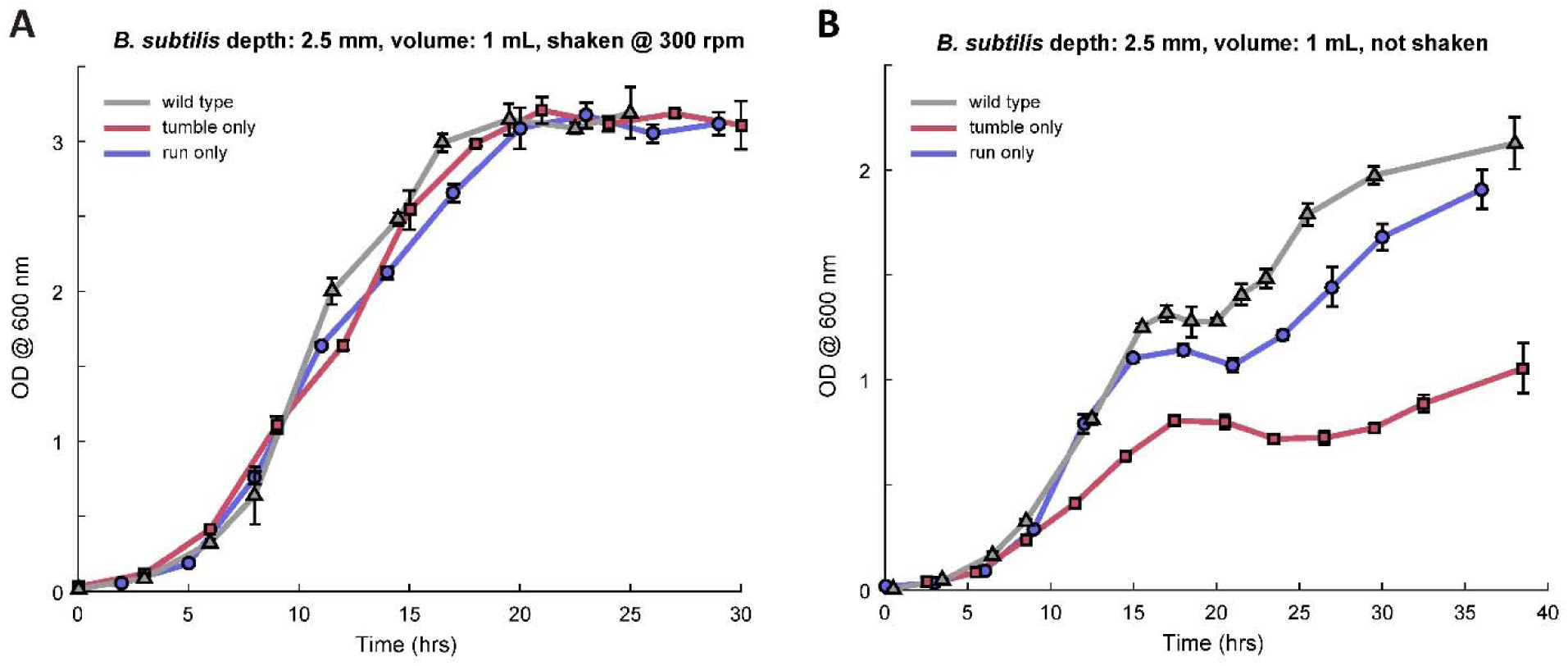
Bioconvection augments growth of wild-type *B. subtilis* over run-only and tumble- only genotypes. Growth curves for *B. subtilis* in 2.5 mm depth of LB media with (A) and without (B) shaking. In media aerated by mechanical shaking (A), the growth rate and carrying capacity for each *B. subtilis* strain were nearly identical. In contrast, growth in media without mechanical shaking (B) showed marked differences in growth rate and carrying capacity between genotypes with wild-type *B. subtilis* possessing the fastest growth rate and greatest carrying capacity, in accordance with its ability to bioconvect. Error bars are standard deviations of the mean over triplicates, and each datum in (B) results from destructive sampling.

Additionally, even though run-only and tumble-only genotypes did not bioconvect, all three genotypes of *B. subtilis* showed transient growth dynamics in the 15–25 hour range that were distinct from the shaken cultures. These transient growth dynamics were most pronounced at media depths of 2.5 and 3.75 mm and were largely absent at media depths of 1.25 and 5.0 mm (SI Fig 1). These transients in OD suggest that chemical and/or physiological differences exist between still and well-mixed conditions that affect bulk growth in similar ways across the three genotypes—independent of bioconvection—and hence may be unrelated to motility or chemotaxis. For instance, one possibility is that some combination of fluid agitation and increased levels of dissolved oxygen prevent biofilm formation in shaken cultures, whereas oxygen stratification and cellular sedimentation in still samples favor a mixed biofilm/planktonic phenotypic strategy that transiently affects measured bulk OD, similar to strategies adopted by other bacterial species (65).

Next, we wanted to know whether *E. coli*, a facultative anaerobe (58), would show similar responses to still vs. shaken environments. We repeated the experiments described above, again using isogenic wild-type, run-only, and tumble-only mutants in the same rich media. Under shaken conditions, the growth dynamics of all three genotypes were statistically indistinguishable (Fig 3A) and approximately sigmoidal (66), whereas unshaken conditions again favored the growth of wild-type over run-only or tumble-only genotypes (Fig 3B). However, genotypic differences in the unshaken growth profiles for *E. coli* were not as pronounced as genotypic differences in the unshaken growth profiles for *B. subtilis*. Given that the Rayleigh number— which controls the onset of convection—is sensitive to the cube of fluid depth, we further hypothesized that the genotype-specific growth rates in an unshaken environment would depend on fluid depth. We again measured the growth curves of all three *E. coli* genotypes in still environments, now at fluid depths of 1.25 mm and 5 mm. All else being equal, these differences in depth corresponded to a factor of 64 in sampled Rayleigh numbers. Fluid depth had a significant, genotypically dependent impact on growth and maximum culture density. At the lowest depth of 1.25 mm (Fig 3B), run-only and tumble-only growth curves were similar, but wild-type displayed a larger growth advantage than what was observed at 2.5 mm (Fig 3C). For the deepest fluid (5 mm), all three genotypes had similar growth curves; however, run-only mutants (which did not bioconvect) had a small but significant growth advantage over wild-type and tumble-only mutants (Fig 3D). We imaged the vertical distribution of cells between these three genotypes (SI Fig 2) and found that run-only cells were more evenly distributed than wild-type or tumble-only mutants, with the latter two accumulating toward the bottom of the fluid column where there is less available oxygen. Taken together, these data show that the growth-rate advantage of bioconvecting wild-type *E. coli* is accentuated at lower fluid depths, whereas shaken cultures display no such genotypic difference.

**Figure 3.**
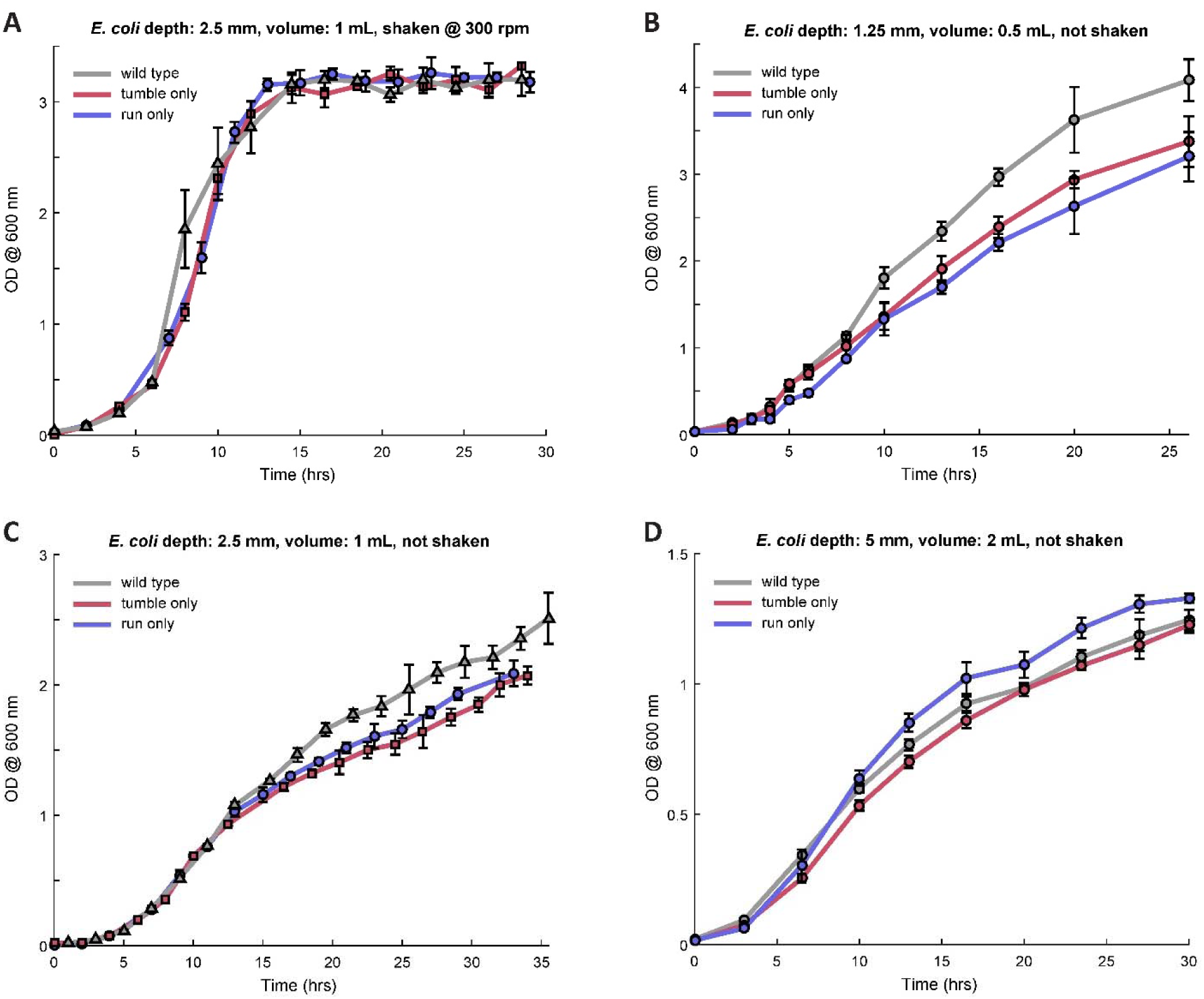
Bioconvection augments growth of wild-type *E. coli* in a depth-dependent manner. Growth curves for *E. coli* genotypes in LB media with (A) and without (B, C, and D) mechanical shaking. The fluid depths were (A, B) 2.5 mm, (C) 1.25 mm, and (D) 5.00 mm. Like *B. subtilis*, at a depth of 2.5 mm bioconvecting wild-type *E. coli* had a growth rate advantage over run-only and tumble-only strains that did not bioconvect in unshaken media, albeit less pronounced than *B. subtilis*. The growth rate advantage of bioconvecting wild-type *E. coli* is augmented in shallower fluid depths (1.25 mm), whereas at 5 mm, the run-only strain reproducibly had a slight growth rate advantage. Error bars are standard deviation of the mean over triplicates, and each datum in (B-D) results from destructive sampling.

These observations highlight a salient difference between the population behaviors of unshaken *B. subtilis* and *E. coli*. In deeper samples, wild-type *E. coli* accumulate toward the bottom of the fluid column where presumably oxygen concentration is lowest, and hence the vast majority of cells cannot contribute to motility-induced mixing (bioconvection or otherwise) that draws oxygen from the air-liquid interface into the bulk of the media. In contrast, *B. subtilis* ascend the oxygen gradient toward the air-liquid interface, thereby creating density inversions that descend through the fluid, aerating it. This difference might explain why *B. subtilis* display a more significant growth-rate dependence on genotype in still cultures as compared to *E. coli* and why *B. subtilis* bioconvective patterns are less sensitive to fluid depth (20, 22).

Finally, the swim dynamics of *E. coli* offer a potential mechanism for why run-only *E. coli* have a slight growth advantage in deeper fluids—relative to wild-type and tumble-only strains— despite not being bioconvective. Run-only mutants cannot respond to chemical gradients (e.g., oxygen) via run-and-tumble chemotaxis, and, unlike *B. subtilis* (67), their swim speed does not depend strongly on oxygen concentration (68, 69). Further, previous work has shown that *individual* bacterial motility augments diffusion (70–72). Thus, run-only mutants do not migrate in any particular direction, and their speed is maintained throughout the fluid column. Consequently, they are more evenly distributed and their concentration near the air-liquid interface is higher as compared to tumble-only mutants or wild-type cells that sediment (SI Fig 2). Thus, we propose that evenly distributed and motile run-only cells augment oxygen diffusion from the air-liquid interface via their individual motility and subsequently increase aerobic respiration and growth. Regardless of the mechanism, these results show that even in the absence of collective motility (bioconvection) or chemotaxis, motility can affect population dynamics.

### Fluid depth and initial cell concentration alter onset and duration of E. coli bioconvection

In all but the deepest still fluid, wild-type cultures of *E. coli* that exhibited bioconvection had a growth advantage over motility mutants that did not bioconvect. This confirmed that these spatially complex and highly dynamic patterns of flow and cellular concentration are linked to increased growth. It also poses the question: which experimental ‘knobs’ can be turned to control the onset, dynamism, and time-course of this advantageous collective behavior? Taking the reductionist view that the Rayleigh number plays a regulatory role in whether the instability that drives bioconvection develops, we sought ways to explore the available parameter space to address this question.

Taking into account a set of physical and biophysical factors, the Rayleigh number controls the transition in transport properties from diffusion to laminar and ultimately turbulent convection. Within its formulation, two such knobs stand out. First, the fluid depth (*L*) can be altered, directly affecting the Rayleigh number as well as bulk oxygen availability via the changing surface area-to-volume ratio. Second, the initial cell concentration can be altered. Previous experiments confirm that for cellular concentrations of OD ∼1 (∼10^9^ cells / mL or volume fraction ∼0.1%), suspensions of rod-shaped bacteria are approximately Newtonian fluids with the same viscosity as the suspending fluid (38). Assuming that near or below these concentrations, the diffusivity of cellular random walks is also independent of cell concentration, we hypothesized that increasing the initial cell concentration would increase the magnitude of mass density fluctuations (Δ*ρ*), thereby increasing the initial Rayleigh number. This increase in initial Rayleigh number should then correlate with reduced bioconvection onset times, because populations either need to grow less to reach the point of bioconvective instability or because cellular suspensions are sufficiently concentrated to initiate the bioconvective instability almost immediately.

We started with the first of these knobs—the depth of the fluid. Similar to the previous data, cultures had an initial OD of 0.05, and we imaged cell-density patterns every 1.5 minutes at fluid depths of 1.25, 2.5, 3.75, and 5 mm. Increasing fluid depth sharply (cubic) increases the Rayleigh number, and thus increased fluid depth should push systems toward turbulent convection and should reduce time to bioconvection onset. First, we visually examined the bioconvective patterns across the four depths at a fixed time point (7 hrs) that coincided with strong bioconvective patterning and exponential bulk growth (Fig 4A, also see SI movies). As fluid depth increased, a visual transition occurred from structured laminar convection to small-scale and irregular turbulent convection (Fig 4A and SI Fig 3B). While the visual signal was clear, we sought a simple and quantitatively consistent way to characterize pattern complexity and timing.

**Figure 4.**
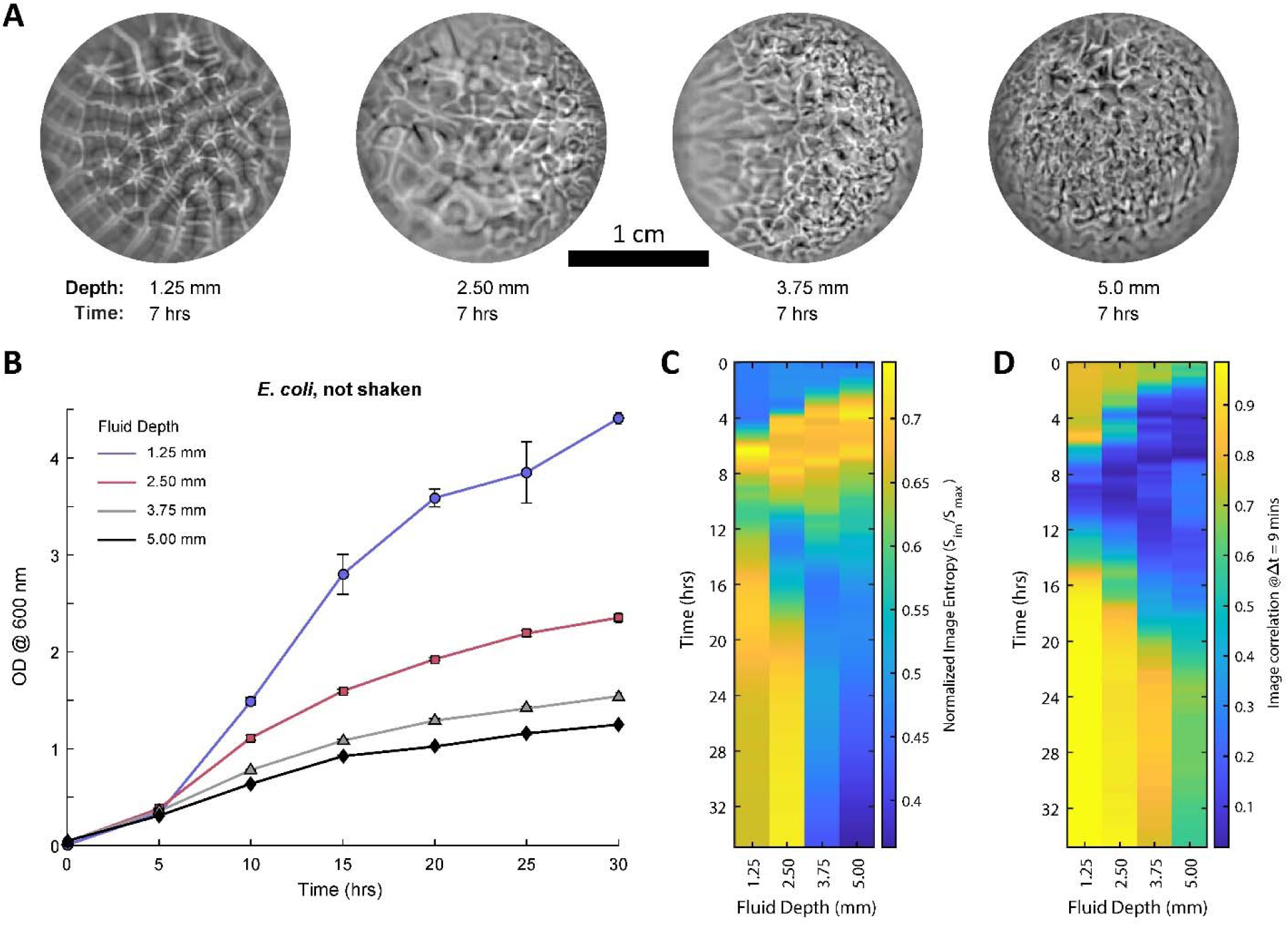
Fluid depth affects the evolution and duration of bioconvective patterns in *E. coli*. Cultures of wild-type *E. coli* were grown in 12-well plates containing LB media without shaking at fluid depths of 1.25, 2.5, 3.75, and 5.0 mm. (A) After 7 hours of growth from an initial density of 0.05 OD, cultures had visually distinct patterns of bioconvection, appearing laminar at the shallowest depth and transitioning to turbulent as depth increased; all depths eventually transiently displayed turbulent bioconvection (see SI Fig 4 movies). (B) Growth curves for unshaken wild-type *E. coli* across the four depths; error bars are standard deviations of the mean over triplicates. (C) Image entropy (a proxy for pattern complexity) as a function of time across the four fluid depths. As fluid depth increased, the time to bioconvection onset decreased, and the number of distinct peaks in image entropy (for t ≲ 10 hrs) increased, indicating emergence of new pattern types with increasing depth. (D) The correlation coefficient between image time points separated by 9 minutes, as a function of time, across the four fluid depths; lower values indicate more rapid change in spatial structure of the pattern; higher values (up to 1) indicate that patterns evolved more slowly. This metric also confirms that onset-time decreases with increasing fluid depth and shows that bioconvection has a longer duration in deeper fluids (approximate duration of the blue region in the barcodes). The stagnation at high correlation values for later times was caused by approximately static cell sedimentation patterns. See SI Fig 4 for full correlation matrices across all replicates. All data are means over triplicates.

To accomplish this, we developed two kinds of time-varying scalar ‘barcodes’ that together gave a simplified and consistent quantification of pattern complexity and temporal evolution, regardless of pattern ‘type’ or the presence or absence of a dominant spatial wavelength. Further, such barcodes needed to be robust to image noise and pixel cropping. We used the scale-invariant image entropy as a measure of pattern complexity for a given moment in time. This measures the information content contained in the whole-image probability distribution of intensities. It does not directly account for spatial structure, but it does account for changes in pattern complexity as represented in the distribution of intensities, which in-turn connects to spatial structure (73). ‘Flatter’ patterns have lower image entropy (a single intensity across the image has zero entropy), and patterns with larger and more dispersed intensity variations have higher image entropy (image entropy is maximized by a uniform random intensity distribution). In Fig 4C, we show the evolution of image entropy across the four fluid depths, averaged in triplicate for each depth. Bioconvective patterns and corresponding entropy barcodes were highly reproducible—see SI Fig 3.

To construct the second barcode, we correlated the cell density for every pair of time points to construct a temporal correlation matrix, whose values indicate how self-similar a pattern is as a function of time (SI Fig 4). Each element in the matrix is a correlation coefficient between −1 and +1 that quantifies the differences in spatial structure and relative intensity values between two time points. Patterns that evolve slowly or have underlying static structure maintain higher correlations from the current time point (i.e., the matrix diagonal), patterns that evolve quickly de-correlate closer to the diagonal. To generate the barcode, for each diagonal time point, we report the correlation value at a fixed future time of 9 mins. This offset was chosen to give high, positive dynamic range to the reported correlation coefficient. Lower values indicate faster pattern evolution, whereas values near 1 indicate that the pattern did not evolve significantly. In Fig 4D, we show how this fixed-future correlation changes through time and across the four fluid depths.

Consistent with the relationship between fluid depth and Rayleigh number, both barcodes showed that increasing fluid depth reduced the time to bioconvection onset—with a factor of 4 increase in fluid depth shifting the onset of bioconvection ∼4 hrs earlier—despite that populations grew faster in shallower fluids (Fig 4B). In addition to our hypothesis, a number of other salient features emerged from these barcodes. First, for otherwise identical cultures, increasing fluid depth increased the duration of bioconvective activity as measured by the correlation barcode (Fig 4D), which is consistent with slower bulk growth at these depths (Fig 4B). Second, increasing fluid depth increased the number of peaks in pattern complexity (i.e., types of patterns) as measured by the entropy barcode and confirmed visually (e.g., compare SI movies 4A and 4D). Third, while ambiguous from the correlation barcodes alone, visual inspection confirmed that the late-time stagnation at relatively high pattern correlations (Fig 4D and SI Fig 4) corresponded to sedimentation patterns that were observed at all depths and in all replicates given enough time (see Fig 4 SI movies for spatio-temporal patterns and eventual sedimentation).

Interrogating the second knob—initial cell concentration—we imaged bioconvection patterns as we varied both the fluid depth (1.25, 2.50, 3.75, and 5.0 mm) and initial cell densities (OD 0.25, 0.5, 0.75, and 1.0) and used those images to calculate the entropy and correlation barcodes, again in triplicate. At a fixed fluid depth, increasing the initial cell density did not significantly alter patterns or their progression (see SI movies for Fig 5). However, consistent with our hypothesis, increasing initial cell concentration shifted the onset of bioconvection to earlier time points, regardless of fluid depth or method of barcode measurement (Fig 5). Additionally, the data showed three other features. First, the degree of time shift was larger at lower fluid depths, primarily due to the fact that in deeper fluids (3.75 mm and 5 mm), bioconvection onset was already nearly immediate. Second, across all fluid depths, increasing the initial cell concentration *decreased* the total duration of bioconvective activity, as measured by the correlation barcode. Given that each experiment starts with the same fixed concentration of nutrients, we speculate that reduced duration with increasing initial cell concentration results from faster consumption of the nutrients that drive motility and hence enable bioconvection. These two observations are further corroborated by the full correlation matrices for this suite of conditions as shown in SI Fig 5. Third, the corresponding time-lapse movies, especially for 5 mm fluid depth, showed fine-scale patterns of bacterial aggregation on the bottom of the dish that may be related to auto-aggregation of high density *E. coli*, as seen in earlier experiments (74).

**Figure 5.**
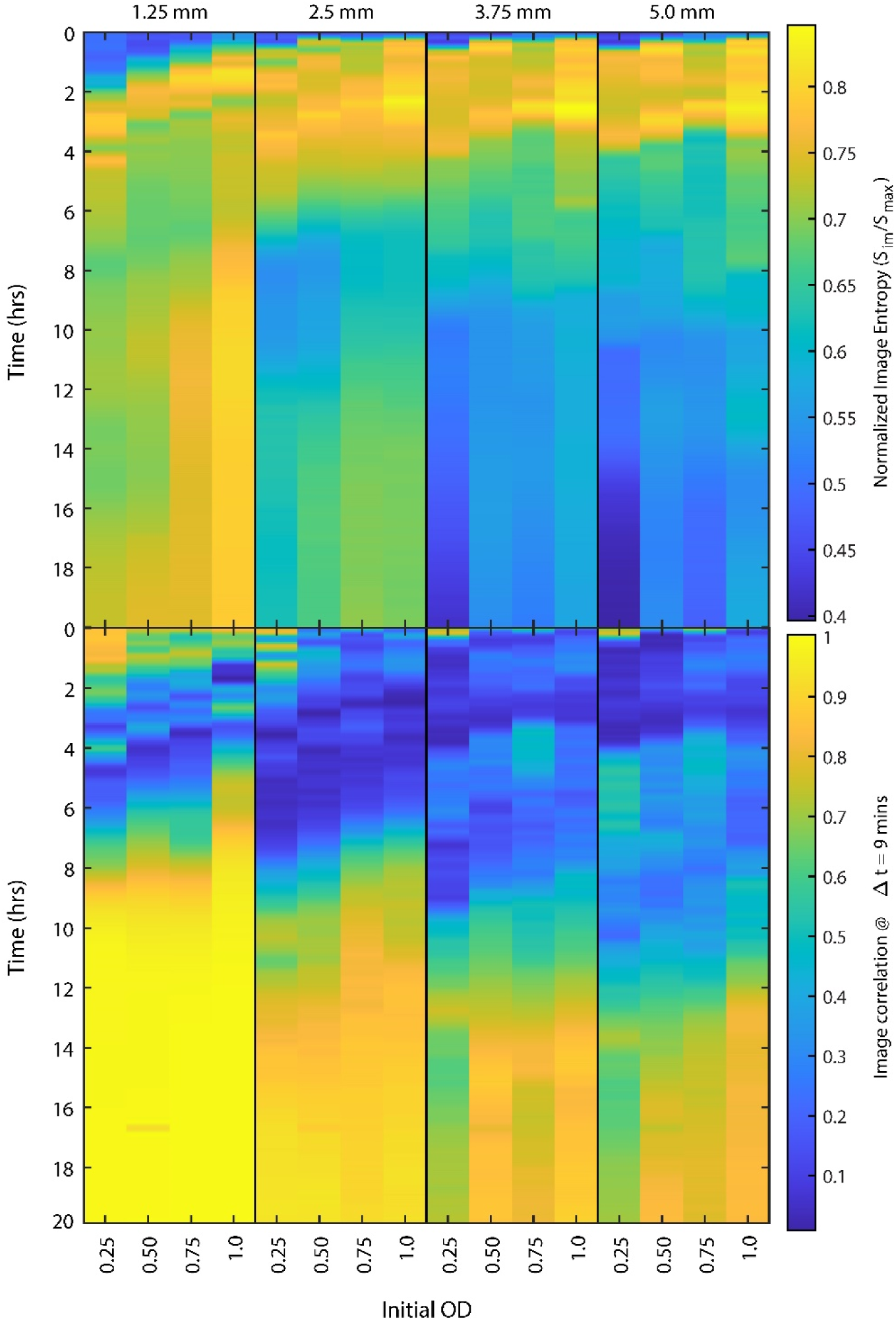
Initial cell concentration and fluid depth have complementary effects on bioconvection timing. To characterize the combined effects of cell concentration and fluid depth, cultures of wild-type *E. coli* with initial OD values of 0.25, 0.5, 0.75, and 1 were placed in 12-well plates with LB media depths of 1.25, 2.5, 3.75, and 5.0 mm. For all 16 conditions, we measured the entropy and correlation barcodes; data averaged in triplicate. Increasing fluid depth and/or increasing initial OD decreased the time to bioconvection onset, though this effect saturates as onset time drops to nearly zero in the deepest and densest samples. The correlation barcodes (bottom) show that increasing fluid depth increased bioconvective duration and increasing initial OD decreased it. The entropy barcodes (top) and visual observation (see corresponding SI movies) showed that initial OD had little effect on the type or progression of bioconvective patterns, whereas, as shown in Fig 4, depth modulated both.

### E. coli bioconvection is robust to changes in the metabolic landscape, oxygen sensing, and carbon-source availability

We found evidence that bioconvection confers selective advantages in certain physical contexts and that physical knobs could alter its timing and duration. Next, we wanted to get a handle on how a few key metabolic inputs affect onset, duration, and patterning of bioconvection. *E. coli* integrate many chemical signals to make the chemotactic decisions that are integral to bioconvection (56, 57, 61). Thus, we did not embark on an exhaustive search to characterize relevant chemotactic inputs. Rather, we sought to make a broad distinction between situations that did or did not offer an initially uniform concentration of readily available carbon (i.e., the sugar galactose). This mandated that we use a synthetic growth medium with control over the available sources of carbon, as well as other vital metabolites like amino acids (see Methods). The fact that *E. coli* exhibit chemotaxis to a wide array of metabolites draws into question whether amino acids alone are sufficient to elicit bioconvection or whether this collective behavior requires the vigorous metabolism and motility that accompany aerobic respiration with readily available carbon.

To answer this, we imaged bioconvection and measured the correlation barcodes for samples across four fluid depths (1.25, 2.5, 3.75, and 5 mm), four initial ODs (0.25, 0.5, 0.75, 1.0), and in media with 0 mM (gal-) or 10 mM (gal+) galactose, again in triplicate; all other media conditions were identical (Fig 6). All of the remaining work characterizes pattern onset and timing using correlation barcodes, as they report directly on pattern dynamics and are less sensitive to background signal than the entropy barcode. A number of familiar trends emerged in these data, regardless of galactose concentration. As in the previous experiments, increasing initial OD shifted bioconvection onset to earlier time points by amounts comparable to previous data in standard rich media (Fig 5). Likewise, increasing fluid depth increased the duration of bioconvection. However, the data also showed a number of distinct trends that depended on the presence of galactose. The duration of bioconvection was significantly reduced in the absence of galactose (Fig 6A, SI Fig 6), and cell-density patterns were distinct between gal+ and gal− conditions (Fig 6B and corresponding SI Movies). Curiously, bioconvection was inhibited at the shallowest depth in gal−, whereas at the same depth, gal+ exhibited bioconvection for the higher initial ODs. Further, in the presence of galactose the high initial ODs at 2.5 mm and all initial ODs at 3.75 and 5 mm had roughly the same duration. These results confirm that *E. coli* bioconvection occurs without readily available carbon but that the presence of galactose significantly increases its duration. The fact that bioconvection was inhibited at the shallowest depths without galactose but occurred at those same depths with galactose is consistent with two, non-mutually exclusive mechanisms. One possibility is that addition of galactose (and the attendant creation of galactose gradients via consumption) alters the run-length of swimming *E. coli* such that the actor-diffusion coefficient *D* decreases, which increases the Rayleigh number. Another possibility is that bacterial density variations are augmented by the addition of galactose; that is, addition of galactose and creation of attendant nutrient gradients alter bacterial motion and cause population-scale movements that increase Δ*ρ*. Both mechanisms relate to motility, which in turn relates to metabolism, and hence the increased duration of bioconvection in gal+ conditions may simply reflect that the presence of readily available carbon (and oxygen) perpetuates motility patterns that increase the Rayleigh number. These two potential mechanisms are difficult to untangle because the cellular diffusion coefficient and the density variations that drive bioconvection are both fundamentally linked to cell motion. Nonetheless, these results demonstrate that, while not required, readily available carbon significantly modulates bioconvection and that the presence or absence of a simple sugar regulates—in a depth-dependent manner—whether a culture will bioconvect. Thus, the growth benefits of adding a metabolite, like sugar, may confer compounding metabolic benefits via bioconvection and in a way that varies with fluid depth.

**Figure 6.**
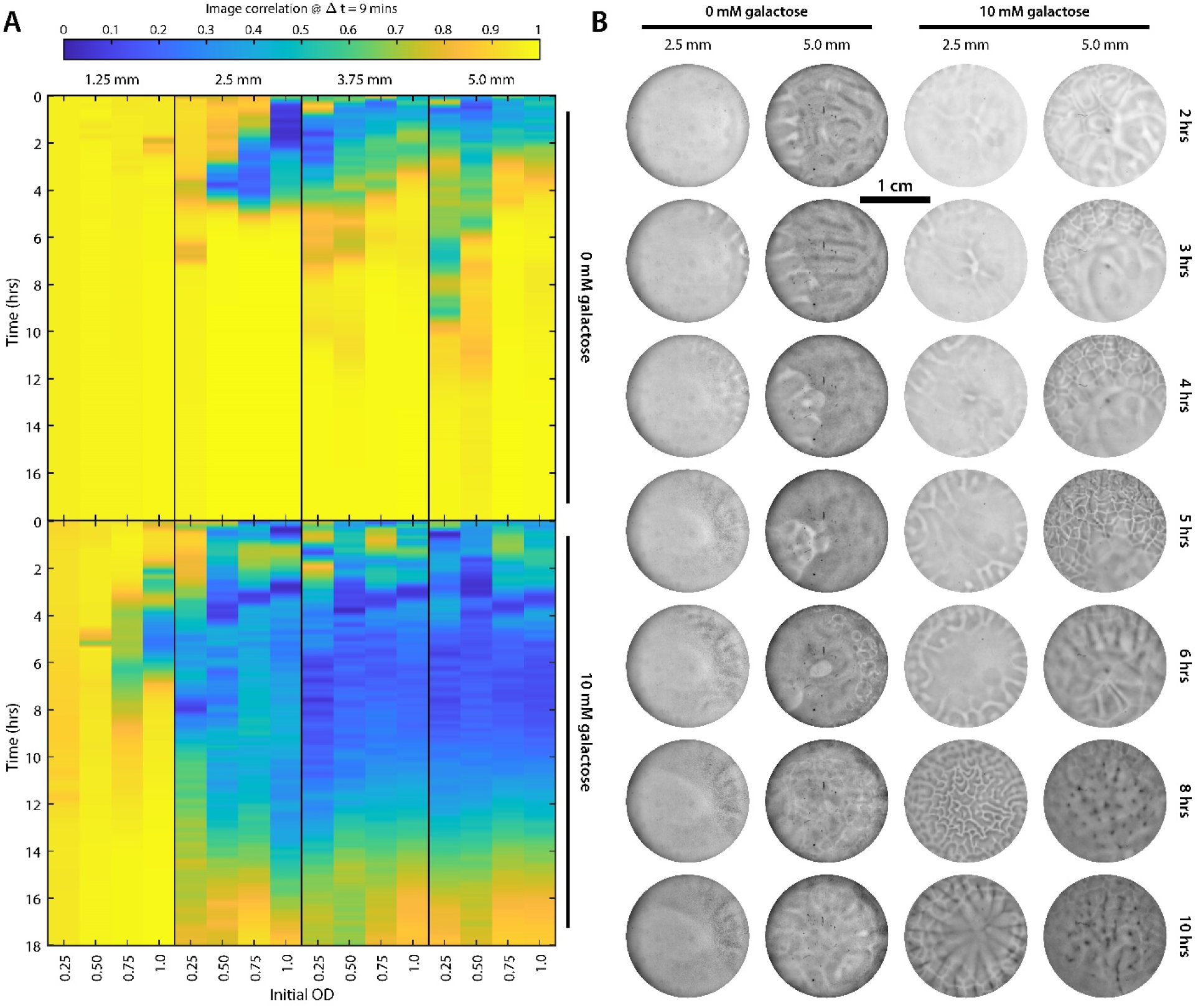
Carbon availability significantly impacts bioconvection. The influence of readily available carbon on bioconvection was explored by growing wild-type *E. coli* in an amino acid-rich synthetic media that either did or did not contain galactose. Like Fig 5, cultures were started at initial ODs of 0.25, 0.5, 0.75, and 1 in fluid depths of 1.25, 2.5, 3.75, and 5.0 mm. (A) The correlation barcodes for scenarios without (top) and with (bottom) galactose show marked differences in onset and timing. Across all but the lowest fluid depth, the presence of 10 mM galactose increased bioconvective duration by a factor of ∼4. At the shallowest depth, bioconvection was inhibited in the absence of galactose. (B) A sample set of images show cell-density patterns at two fluid depths, across time for conditions with and without galactose starting from an OD of 0.25. Full movies for each column of (B) are available in the SI.

Next, we wanted to understand if metabolite sensing—specifically oxygen sensing or energy sensing—is required for bioconvection and/or if its removal had significant effects on onset, duration, and patterning of bioconvection. In order to determine whether aerotaxis is required for bioconvection, we imaged mutant *E. coli* cultures unable perform aerotaxis (Δ*aer* mutants (75)). Mutant strains were grown in varying fluid depths (1.25, 2.5, 3.75, and 5.0 mm) from an initial OD of 0.5 (Fig 7, SI Fig 7). The aerotaxis-null strains not only bioconvected but exhibited the same trends in onset timing and duration with fluid depth that were observed in wild-type cultures (Fig 5). Further, as assessed visually, the bioconvective patterns exhibited by the aerotaxis-null strains were similar in form and progression to those seen in wild-type cultures at the same depths, with the notable difference of increased sedimentation in the aerotaxis-null strains. These results not only indicate that bioconvection can occur without direct oxygen sensing (unlike in *B. subtilis*), but the similarly to wild-type in other measured aspects suggests that direct oxygen sensing is not a significant input to wild-type *E. coli* bioconvection.

**Figure 7.**
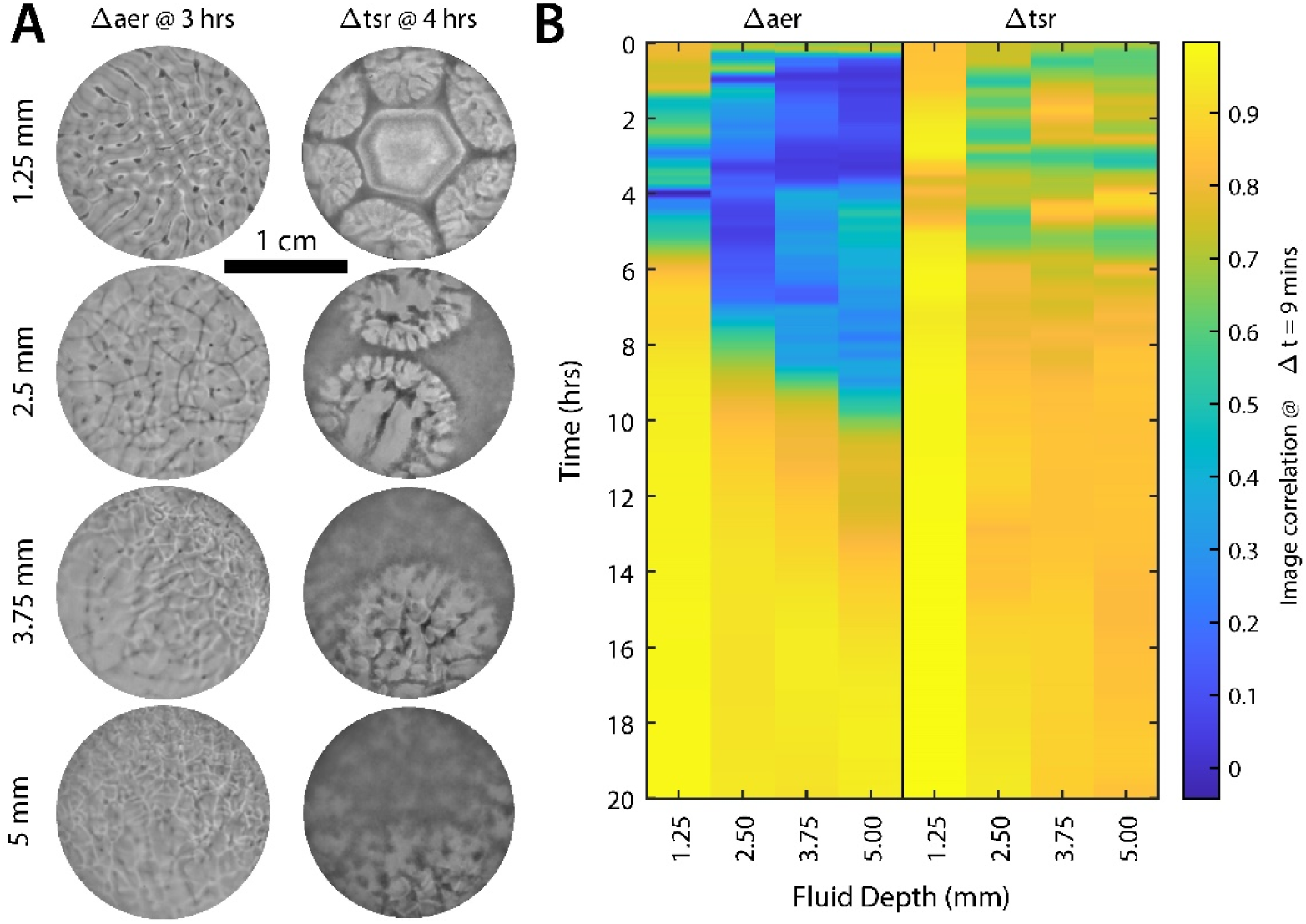
Neither aerotaxis nor ‘energy-taxis’ are required for *E. coli* bioconvection. To determine if direct or indirect oxygen sensing is required for bioconvection in *E. coli*, aerotaxis- deficient (Δ*aer*) and energy-taxis-deficient (Δ*tsr*) strains were imaged for the presence of bioconvective patterns. Both mutant strains of *E. coli* were initialized at OD 0.5 and placed into 12-well plates at fluid depths of 1.25, 2.5, 3.75, and 5.0 mm. (A) Images of cell-density patterns in *E. coli* mutants unable to directly sense oxygen (aerotaxis-deficient) (left column) or unable to perform ‘energy-taxis’ (Δ*tsr*) (right column). (B) Corresponding correlation barcodes for aerotaxis and energy-taxis-deficient mutants at four fluid depths; each barcode is an average over three experiments. Imaging data (including corresponding SI movies) and barcodes show that both mutants can bioconvect at all fluid depths. In aerotaxis-deficient mutants, cell-density patterns and trends in onset and timing are similar to wild-type (e.g., compare with Fig 5 at 0.5 OD), indicating that direct oxygen sensing is not a major driver of *E. coli* bioconvection. Energy-taxis- deficient mutants, however, display distinct patterns and high levels of cell sedimentation, indicating significant disruption of wild-type bioconvective behavior.

Oxygen gradients can directly drive collective bacterial movements (as shown in *B. subtilis*), but oxygen can also *indirectly* create metabolic cues that affect migration. Oxygen is not only a potent chemoattractant in some species but is also an electron acceptor in oxidative phosphorylation, which allows cells to generate more ATP from available nutrients than during anaerobic respiration (57, 76). Therefore, the presence of oxygen spurs rapid metabolic activity with corresponding increases in the rate of local nutrient consumption (58); this creates local nutrient gradients (the cells are sinks), which in turn spur chemotaxis in response to those gradients. Such mechanisms are known to create collective and persistent migration of cells in low Rayleigh number systems that do not bioconvect but instead exhibit bacterial waves (53, 54, 77). Thus, even in the absence of direct oxygen sensing, consumption itself can create metabolic cues for migration.

Energy-taxis is one such form of chemotaxis that can respond to metabolite gradients produced through local consumption. It occurs when cells tumble and reorient in response to a decrease in internal energy levels as sensed by their transmembrane proton gradient (26, 78)— thus energy-taxis is a non-specific response to current metabolic conditions. We reasoned that if bioconvection was abolished or significantly disrupted in the absence of energy-taxis that would support the hypothesis that *E. coli* use local gradients resulting from consumption to create the density variations that drive bioconvection. To explore this mechanism, we examined bioconvection in energy-taxis-deficient *E. coli* mutants (Δ*tsr* mutants (78)). Mutant strains were grown in varying fluid depths (1.25, 2.5, 3.75, and 5.0 mm) from an initial OD of 0.5 (Fig 7, SI Fig 7). The energy-taxis-deficient cultures did bioconvect; however, the patterns, as well as their progression through time, were distinct from wild-type (see SI movies for Fig 7). Additionally, as compared to all other conditions, sedimentation was severe, and, correspondingly, the correlation barcodes had low dynamic range (Fig 7B). Further, we observed that bioconvective flow had the effect of patterning cells sedimented on the bottom surface (see SI movies). While this significant disruption of bioconvection is consistent with the hypothesis that local gradients drive *E. coli* bioconvection, it is difficult to draw stronger conclusions via a comparison to wild-type behavior, as Δ*tsr E. coli* are also deficient in an array of other behaviors, including serine sensing (79) and auto-inducer-2-based quorum sensing (80). Thus, alternatively, Δ*tsr*-mediated changes to bioconvection may indicate that auto-inducer-2-based quorum sensing is a significant part of the chemotactic response that elicits bioconvection in *E. coli*.

Finally, the chemotactic and metabolic preferences of *E. coli* (56, 57), as well as their ability to bioconvect without direct oxygen sensing, led us to question whether they could bioconvect without an oxygen gradient that emanates from the air-liquid interface. We measured the presence, onset time, and duration of bioconvection in samples *without* an air-liquid interface. We removed the interface by sealing cultures of wild-type *E. coli* between two glass plates separated by a flat Teflon ring (thicknesses of 0.5, 0.8, 1, 1.5, 2.3, 3.2, and 4.8 mm, all with the same diameter as a 12-well plate). This abolished exogenous oxygen flux and the corresponding interfacial gradient. It also removed differences in growth rate resulting from changes in surface area-to-volume ratio and corresponding variations in bulk oxygen availability. Sealing the chamber removed the meniscus, stabilized the fluid layer (e.g., prevented beading), and drastically reduced evaporation, such that we could now sample fluid depths lower than 1 mm, which we could not sample in open air. Cultures were grown and time-lapse imaged in the same rich defined medium used in the galactose experiment. We note that the correlation barcode cannot distinguish between bioconvection and the related phenomenon of bacterial waves (54, 55, 77), which are, like bioconvection, transient and coherent fluctuations in bacterial density due to their chemotactic and motile activity.

In Fig 8A, we show correlation barcodes across four distinct initial ODs and seven distinct fluid depths. These data, in combination with confirmatory imaging (Fig 8B and corresponding SI movies), showed that *E. coli* bioconvection does occur without an air-liquid interface. Like experiments with an air-liquid interface, samples that lacked an interface displayed bioconvective patterns earlier with increased initial OD, and increasing the fluid depth increased the duration of bioconvection. However, as compared to samples with an air-liquid interface, samples lacking an interface showed reduced duration of bioconvective activity as measured by the correlation barcodes. Below 2.3 mm fluid depth, and across all initial ODs, bioconvection lacking an interface was short-lived, lasting at most an hour, whereas samples with an interface displayed bioconvection for ∼6 or more hours across all heights and initial ODs (Fig 5). At the lowest depth and initial OD—where the Rayleigh number is smallest—bioconvection was absent. Curiously, while this set of experiments sampled the lowest fluid depths, with correspondingly lower Rayleigh numbers, our imaging did not display patterns of laminar convection; bioconvection resulted only in turbulent convection. Lastly, for *t* ≥ 8 hrs, bubbles reproducibly formed in the sealed media volume, especially at lower fluid depths, likely due to the build-up of gaseous metabolic byproducts (e.g., CO_2_) that could not exit through an air-liquid interface. All pixels corresponding to bubbles were removed in further analysis via segmentation (see Methods). While it is not clear how, in the absence of an air-liquid interface, a cue for vertical ascent is generated, one potential origin is that the combination of sedimentation and consumption in proportion to cellular concentration sets up a gradient that, on average, vertically orients migration and thus elicits bioconvection.

**Figure 8.**
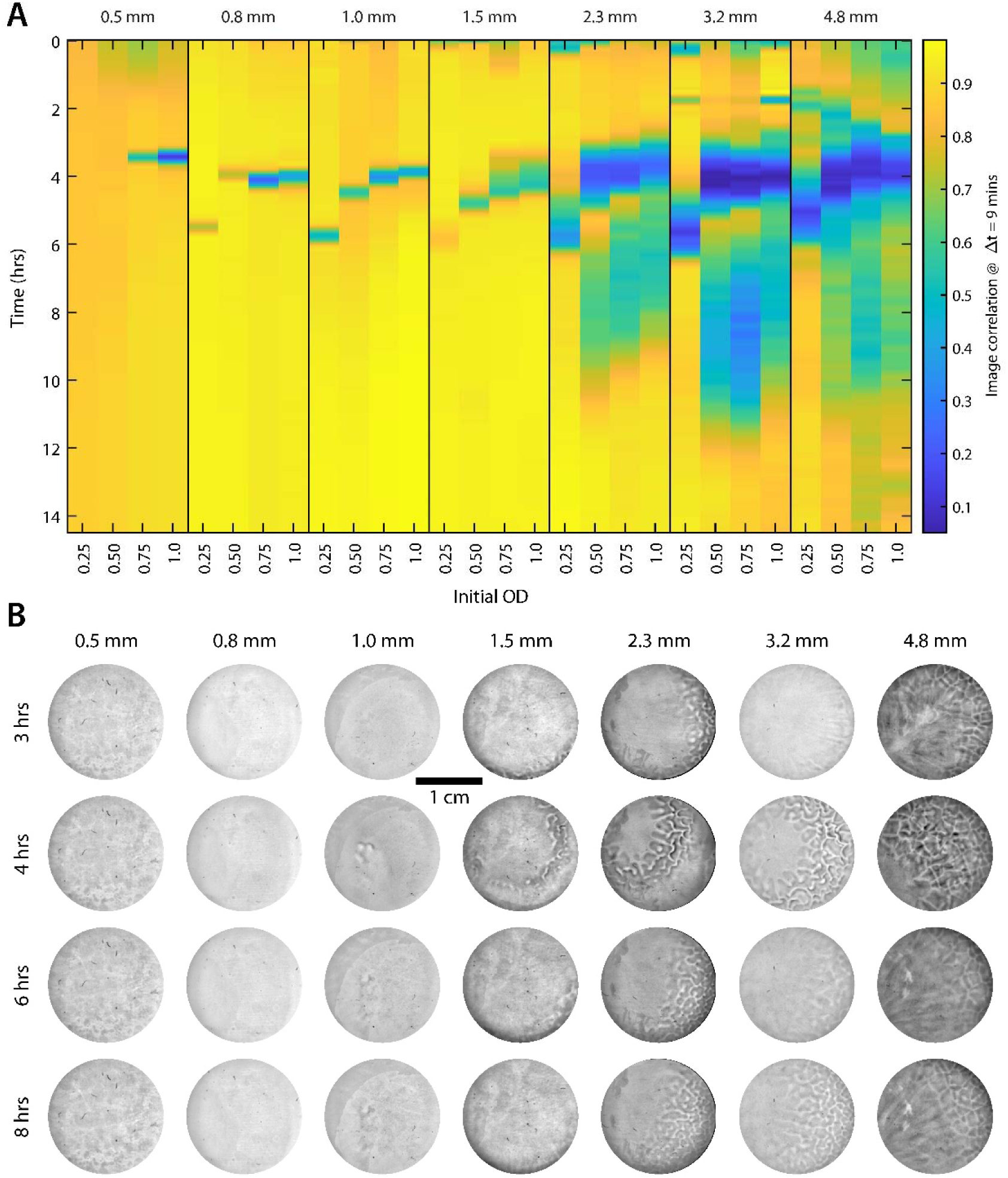
*E. coli* do not require an air-liquid interface to bioconvect. To determine if *E.* coli require an air-liquid interface to bioconvect, we grew wild-type cultures in rich defined media with galactose in sealed, air-tight chambers across multiple fluid depths and initial ODs. (A) In all but the shallowest depth, bioconvection onset time decreased with initial OD. Above 1.5 mm, bioconvection exhibited the familiar trends of increasing in duration with fluid depth and decreasing in duration with increasing initial OD. Below 2.3 mm, samples typically exhibited a ‘pulse’ of bioconvection lasting an hour or less. Bioconvection was completely inhibited at the lowest initial ODs and shallowest depth. All barcodes are means of experiments performed in triplicate. (B) Representative images of cell density across all depths for initial OD 0.75. Interestingly, despite sampling lower fluid depths with correspondingly lower Rayleigh numbers, we did not observe laminar convection in these samples.

Together with the previous data that oxygen sensing is neither required for nor has significant effects on bioconvection, that energy sensing *does* have a significant effect on bioconvection, and that carbon source availability under aerobic conditions affects duration, we conclude that direct oxygen sensing is not a key player in *E. coli* bioconvection; instead, consumption-generated gradients, catalyzed by aerobic metabolism, are the primary drivers of *E. coli* bioconvection. Therefore, *E. coli* may be a far more versatile bioconvector, able to migrate and bioconvect in response to a wider set of metabolic cues. In addition to differentiating *E. coli* from other canonical bioconvectors, these results are salient because the ability to bioconvect in the absence of an air-liquid interface and its attendant gradient, in the absence of key, genetically encoded chemical-sensing mechanisms or in the absence of a readily available carbon source, greatly expand the set of the contexts in which microbial bioconvection might be expected.

## Discussion

Bioconvection is a complex phenomenon that combines individual cellular behaviors (chemical sensing, motility, chemotaxis, and metabolism), collective motion, and the spatial gradients of multiple chemical species, with fluid properties and dynamics to produce large-scale flow and group-level selective pressures. By imaging the cellular density field and measuring bulk OD, we found that bioconvection could, within certain physical contexts, confer a growth benefit relative to otherwise metabolically identical strains that did not bioconvect in the same physical context. Further, we found that turning the salient knobs of fluid depth and initial cell concentration altered the Rayleigh number that controls the onset of bioconvection. It is worth noting that the full complexity of the experimental system is captured by the dynamic, three dimensional scalar fields of cellular concentration and multiple chemical species, as well as the co-evolving vector flow field; these are richer data that could be used to ask more precise biological questions and make deeper connections with physical theory. However, methods to simultaneously and non-invasively measure all of these features do not yet exist, and the challenges of analyzing such enormous data sets would be substantial. Nevertheless, our experiments confirm that bioconvection presents a context in which population-level migration via chemotaxis can be advantageous due to the physical effects of mixing, independent of the *individual* metabolic benefits of ascending a particular chemical gradient.

Along similar lines, we showed in this work that the presence of a simple metabolite (here galactose) could regulate the occurrence, onset, and duration of bioconvection in a depth-dependent manner. We find this particularly interesting because (i) it shows that the concentration of a chemical species can influence the physical Rayleigh number through the actions of cellular populations (e.g., increased motility or chemotactic response); (ii) it suggests that selective pressures applied to a population by the physics of bioconvection (Figs 2 & 3) can be modulated by chemical inputs; (iii) it indicates that for fluid systems without external agitation, the presence of a metabolite can have compounding effects on the population—both impacting metabolism directly (as through respiration) and by spurring large-scale flow that affects mixing and distribution of other metabolites (e.g., oxygen) (81); and (iv) it shows that through the action of cellular populations, addition or depletion of a chemical component can modulate large-scale flow in the environment as a whole, with implications not only for those cells but any other organisms residing in the fluid column.

It is also clear from our data that fluid depth and oxygen play complicated and intertwined roles in eliciting *E. coli* bioconvection. In the presence of an interface, changes in fluid depth entangle changes in Rayleigh number with changes in the surface area-to-volume ratio, the latter directly affecting bulk oxygen availability. As confirmed in SI Fig 1, in deeper un-agitated fluids, wild-type cells grow more slowly, presumably due to reduced oxygen availability per unit volume. Given the fixed initial concentration of nutrients in the media (not including oxygen), the reduction in growth rate with fluid depth indicates an inverse relationship between the per-volume rate of aerobic metabolism and the duration of bioconvection. The aerotaxis-deficient mutants also exhibited the same trends. It is not surprising then that removal of a readily available carbon source (galactose) reduces the nutrient pool for aerobic respiration and thus reduces the duration of bioconvection. These data suggest that, at least in *E. coli*, the duration of bioconvection is regulated by metabolism within a fixed pool of non-oxygen metabolites. However, we observed that depth also increased duration in samples that did *not* have an air-liquid interface. These scenarios lack exogenous oxygen flux, and thus levels of carbon and oxygen were fixed. These results indicate that the metabolism that drives motility (and ensuing bioconvection) has a depth dependence even without changing the surface area-to-volume ratio. A potential mechanism consistent with the metabolism vs. duration hypothesis is that cell sedimentation, which itself depends on fluid depth, causes a differential rate of metabolism along the fluid column. Cells collect near the bottom and locally deplete nutrients, and the corresponding consumption gradient orients cells for vertical ascent via chemotaxis, driving bioconvection. Simultaneously, the upper portion of the fluid retains nutrients longer than if the same number of cells were isotropically distributed within the fluid column, and thus duration of bioconvection is extended as fluid depth increases. In any case, the fact that fluid depth strongly modulates bioconvection means that depth impacts active nutrient mixing and the dispersal of chemical signals (81, 82)—and hence alters the selective context in which a population grows. Thus, across multiple physical, metabolic, and genetic contexts, the depth of an un-agitated fluid can be both a potent modulator of growth and a selector for particular cellular behaviors via bioconvection.

Finally, the fact that bioconvection is a distinctly population-level behavior that connects metabolism, motility, and growth rate to large-scale fluid motion and attendant changes in metabolite concentration suggests that as much as individual and collective behaviors determine whether a population bioconvects, bioconvection has the potential to shape behavior. For example, *E. coli* are motile under a wide array of metabolic conditions, and recent work has shown that investment in motility (which is required for bioconvection) is in proportion to the growth benefits that it confers (83). A particularly intriguing possibility is that—all else being equal—bioconvection does not necessarily select for identical individual behaviors. It is not yet known whether populations under bioconvective conditions benefit from a mixture of phenotypic and/or genotypic behaviors relative to isogenic populations (like those explored here) or whether this dynamic environment opens niches for other species and/or behaviors with wider ecological effects. For instance, one population of cells might consume a particular metabolite and thereby produce gradients that other cells follow to produce density variations that drive bioconvection. Likewise, in many biological contexts, populations can (genotypically or phenotypically) split into ‘cooperators’ and ‘cheaters’ (84–86), with the former producing some kind of ‘public good’ that is exploited by the latter. We speculate that during bioconvection, large-scale fluid motion *is* the public good because such flows disperse metabolites (e.g., oxygen) that provide utility for cheaters and cooperators alike. It remains to be seen, then, how the dynamics of mixed populations composed of cooperators and cheaters evolve under physical and metabolic conditions conducive to bioconvection and, moreover, how the physical attributes that modulate bioconvection, like fluid depth or viscosity, alter the balance between microbial sub- populations exhibiting distinct behaviors.

## Methods

### Bacterial Strains

Wild-type, run-only (Δ*cheB*), and tumble-only (Δ*cheY*) *B. subtilis* 3610 strains were generously provided by the lab of Prof. Daniel Kearns (Indiana University). Wild-type MG1655 *E. coli* were generously provided by Prof. KC Huang (Stanford University). The run-only (Δ*cheW*), tumble-only (Δ*cheY*), aerotaxis-deficient (Δ*aer*), and energy-taxis-deficient (Δt*sr*) strains of *E. coli* were part of the KEIO collection (87) and were purchased from the *E. coli* Genetic Stock Center.

### Growth Media and Culturing Bacteria

Bacteria were stored at −80 C in aliquots of glycerol stocks prepared by mixing 50% glycerol 1:1 with saturated overnight cultures of individual strains grown in Lysogeny Broth (LB) (Lennox formulation). Prior to experimental setup, bacterial cultures were prepared with a 1:1000 dilution of a frozen glycerol stock into either LB (Lennox formulation) or a defined rich media composed of 1mM magnesium sulfate, 1 mM ammonium sulfate, 35 mM sodium chloride, 20 mM potassium phosphate (pH 7.4), 500 uM calcium chloride, 50 nM ferric nitrate, 10 uM thiamine, 10 mM galactose, 1x MEM amino acids solution (ThermoFisher), 1x MEM non-essential amino acids solution (ThermoFisher), and 2 mM L-glutamine. Cultures were grown shaking at 200 rpm at 37 C. After reaching exponential phase at OD_600 nm_ = 0.5, cultures were spun down at 1,500 *g* for 10 minutes and resuspended in an equal volume of fresh growth media; this spin down and re-suspension was repeated a total of three times in order to replace lost nutrients and remove metabolic byproducts. Cultures were then diluted appropriately for each experiment.

### Experimental Preparation

Bacterial cultures were placed into either costar 12-well plates (Corning) open to air or sealable wells (no interface) custom made from glass and Teflon. Custom chambers were created by drilling 22.1 mm diameter holes, in the exact arrangement of a 12- well plate, into sheets of Teflon (ePlastics Inc.) with varying thicknesses. Teflon sheets were then lightly coated in inert vacuum grease (Dow Chemicals Inc.) before being placed onto a 2.5 mm thick glass sheet. Cultures were then placed in each Teflon well before being sealed with a second glass plate without bubbles.

Samples were protected from ambient and microscope-originating thermal variations by enclosing them in a glass-windowed insulating chamber made from plastic and Styrofoam. A clear glass heating plate (Thermo Plate, Tokai Hit Inc.) was placed on top of the insulating chamber and set to room temperature to act as an active thermal buffer during experimentation. We imaged fluorescent beads over time to verify that identically prepared samples lacking bacteria did not exhibit bulk flow (though beads do move due to thermal diffusion). In samples with an air-liquid interface, 12-well plate covers were coated in an anti-fogging compound (Rain-X) to prevent condensation during experiments. Plates were mounted on an automated and motorized stage and imaged with a Nikon SMZ25 dissection scope equipped with a 2X SHR Plan Apo lens. Images were taken with a Zyla 5.5 CMOS camera (Andor Inc.) at regular intervals (3 minutes in Fig 1; 1.5 minutes in all other experiments) over 12 to 32 hours as indicated in the figures.

### Image Analysis

Images were cropped to the approximately circular region of each well and any imaging abnormalities (e.g., bubbles in the no-interface experiments, aberrations near the edge of the well) were removed by image masking with custom Matlab (The Mathworks, Inc.) scripts. Typical image sizes were ∼900 x 900 pixels with 20.26 um/pixel. Images were then corrected for non-uniform background illumination using a Fourier band-pass filter in FIJI (NIH) (2 to 150 pixel wavelength passing) with mask correction. Filtered images were then contrast adjusted with custom Matlab scripts to remove 0.001% outliers from the whole-stack histogram of intensities; thus, image intensities are comparable across each stack. All movies and figures show these illumination- and contrast-adjusted images. To reduce the influence of edge effects, an annulus of ∼150 pixels was removed from the outer circular edge of each image before calculating barcodes. The entropy of each image was calculated by taking these post-processed pixels, calculating the probability distribution of intensities on an 8-bit scale, and then calculating the normalized entropy 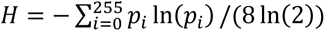. Correlations between pairs of images were found by calculating the mean-subtracted, norm-squared vector representation for each set of post-processed pixels in an image and then calculating the dot-product of those vectors between every pair of images—those values form the correlation matrices in the SI figures. The correlation barcodes are then the values of the correlation matrices offset from the diagonal by the fixed time of 9 minutes. All of these procedures were performed using custom Matlab scripts that are available on the publisher website as SI material. All growth plots, barcodes, and matrices shown in the figures are averages across triplicates for each condition.

## Acknowledgments

We thank Prof. Dan Kearns for providing the *B. subtilis* strains, Prof. KC Huang for providing the wild-type *E. coli*, and the University of Oregon for funding this research. DS collected all data; DS and TU analyzed data, constructed figures, and wrote the paper.

## Supplemental Figures

**SI Figure 1.**
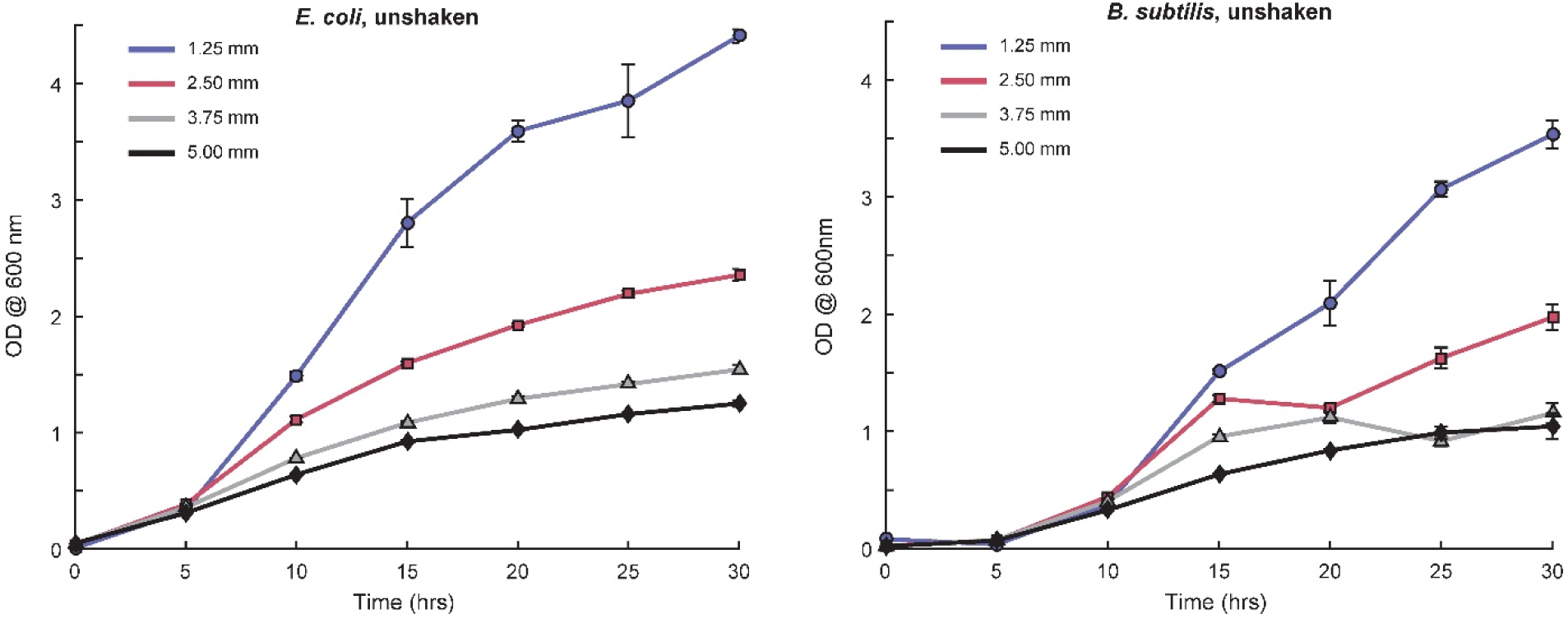
Growth rates of wild-type *B. subtilis* and *E. coli* are depth-dependent. Lower time- resolution growth curves for bioconvecting wild-type *B. subtilis* and *E. coli*. Decreasing fluid depth increased the surface area-to-volume ratio, reduced the Rayleigh number, and robustly sped up growth in both species. Error bars are standard deviation of the mean over triplicates.

**SI Figure 2.**
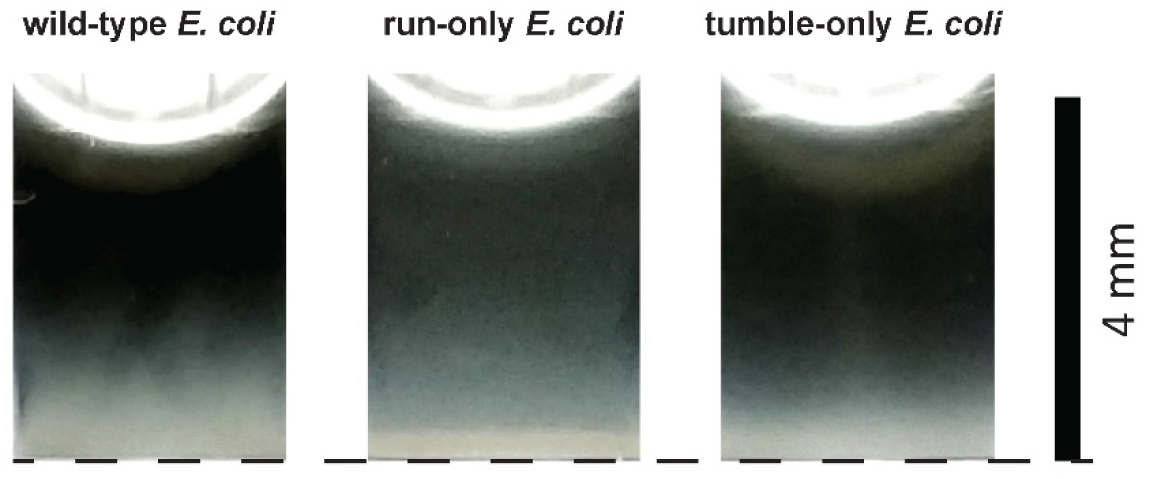
In deeper fluids, run-only *E. coli* are more evenly distributed in the fluid column than wild-type or tumble-only strains. Wild-type, run-only, and tumble-only strains of *E. coli* were grown in LB media under bioconvective conditions to an OD of 0.5. 200 µL of each culture were placed into a still cuvette (∼4 mm depth) for 30 mins, at which point the vertical distribution of cells was imaged via dark-field microscopy. At this depth, wild-type and tumble-only strains showed notable sedimentation, whereas run-only mutants were more evenly distributed throughout the fluid column.

**SI Figure 3.**
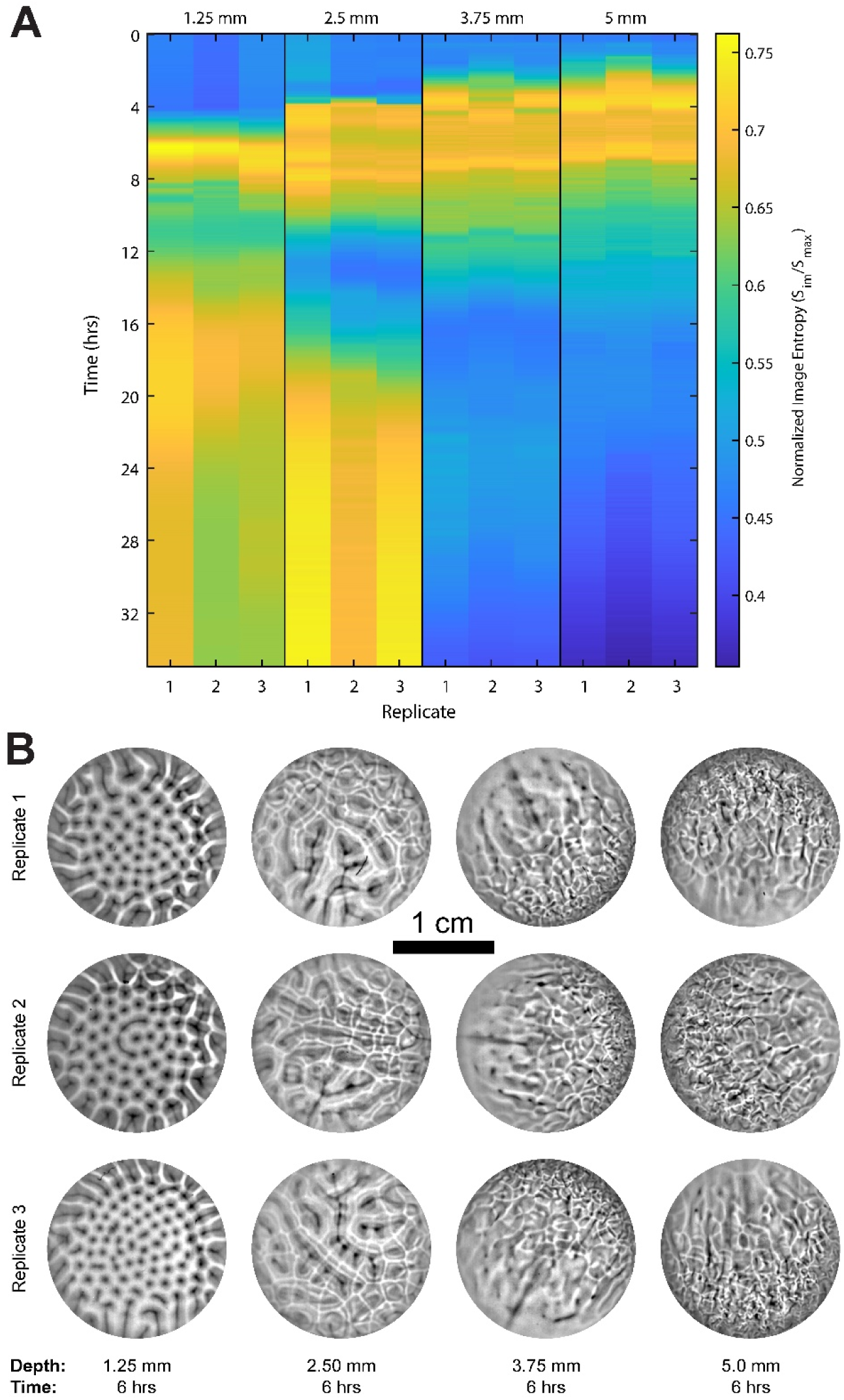
Reproducible evolution of bioconvective patterns and barcodes at various fluid depths. Cultures of wild-type *E. coli* were grown without shaking from a starting OD of 0.05 at fluid depths of 1.25, 2.5, 3.75, and 5.0 mm in 12-well plates. (A) Each pane shows three replicates of the temporal evolution of image entropy for cells grown at different fluid depths. For each depth, the time course of image entropy showed consistent trends between replicates. (B) At each fluid depth, bioconvection produced remarkably similar patterns of cell density across identical replicates. Progressing from shallower to deeper fluids, those patterns displayed reproducible transitions from laminar to turbulent convection.

**SI Figure 4.**
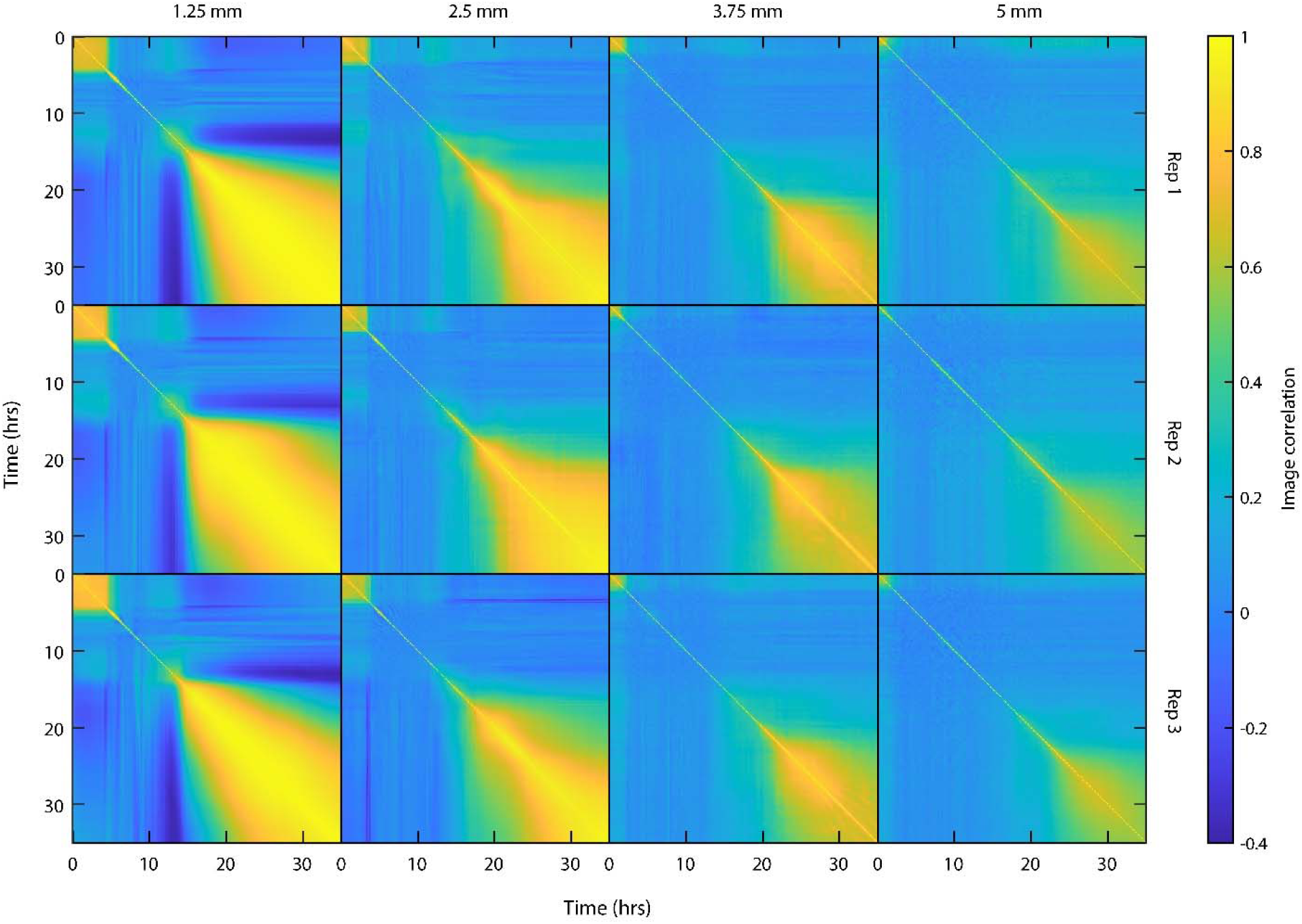
Cell-density correlations across fluid depth, time, and replicate. Each pane shows the cell-density correlation matrix for a particular fluid depth and replicate. In each matrix, an element (colored pixel) quantifies the spatial and intensity correlations between a given time point located on the diagonal and a future (right) or past (left) time point. Increasing fluid depth: (i) reduces the onset time (the ‘yellow squares’ in the upper-left of each pane), (ii) increases the duration of bioconvective dynamics (from the lower-right corner of the yellow squares to the larger yellow regions in the lower-right corner of each matrix), and (iii) at later times all samples displayed sedimentation that produced the larger regions of consistently higher correlation.

**SI Figure 5.**
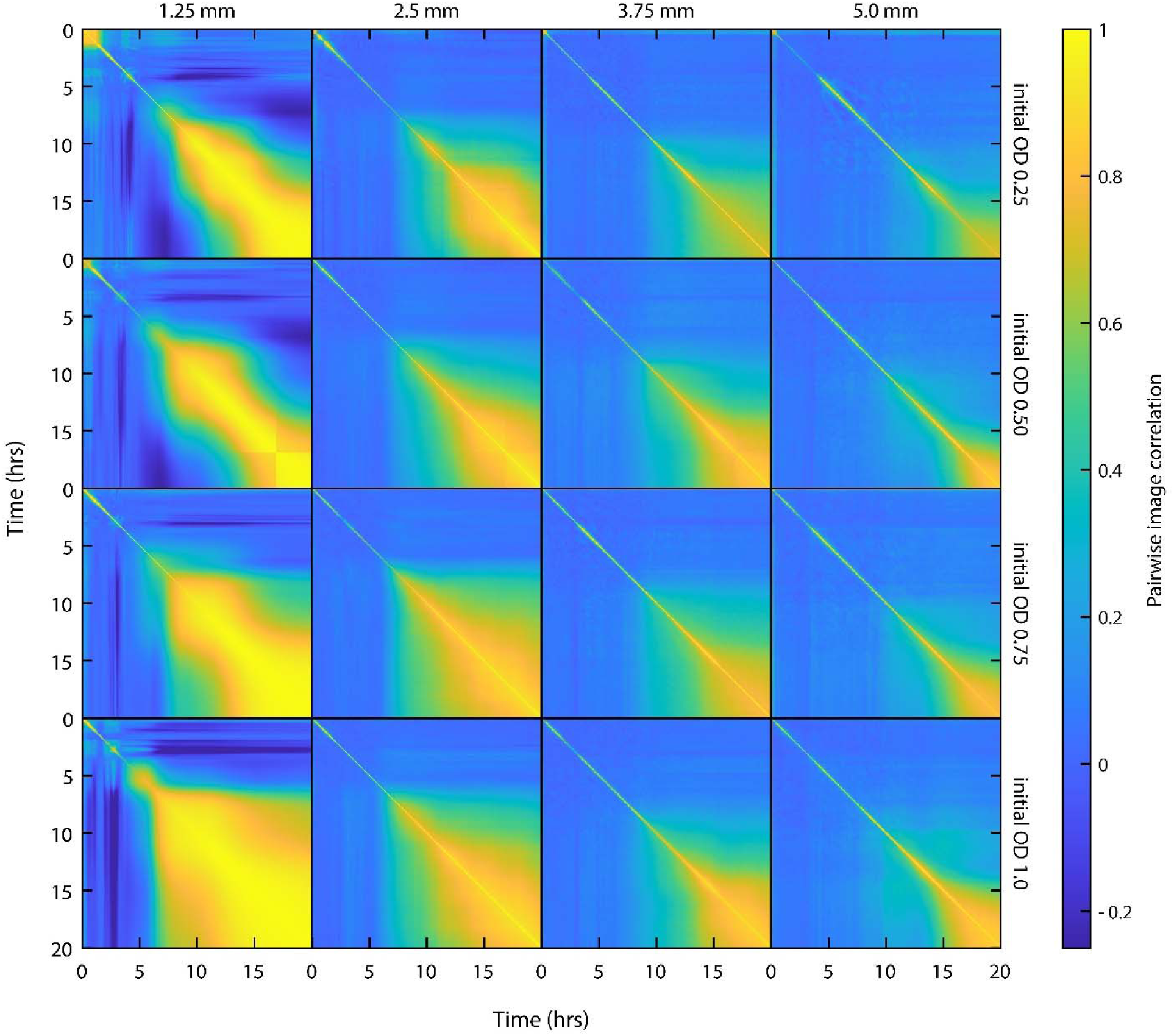
Cell-density correlation matrices across fluid depths and initial cell concentrations. Each pane shows the full correlation matrix of cell-density (image) patterns for each fluid depth and initial OD; each pane is an average over three experiments. These data give a richer view of how patterns evolve in time under the changing experimental conditions. The matrices corroborate the conclusions drawn from Fig 5 and confirm that, as measured by the correlations, changes in pattern evolution were more sensitive to fluid depth than initial OD.

**SI Figure 6.**
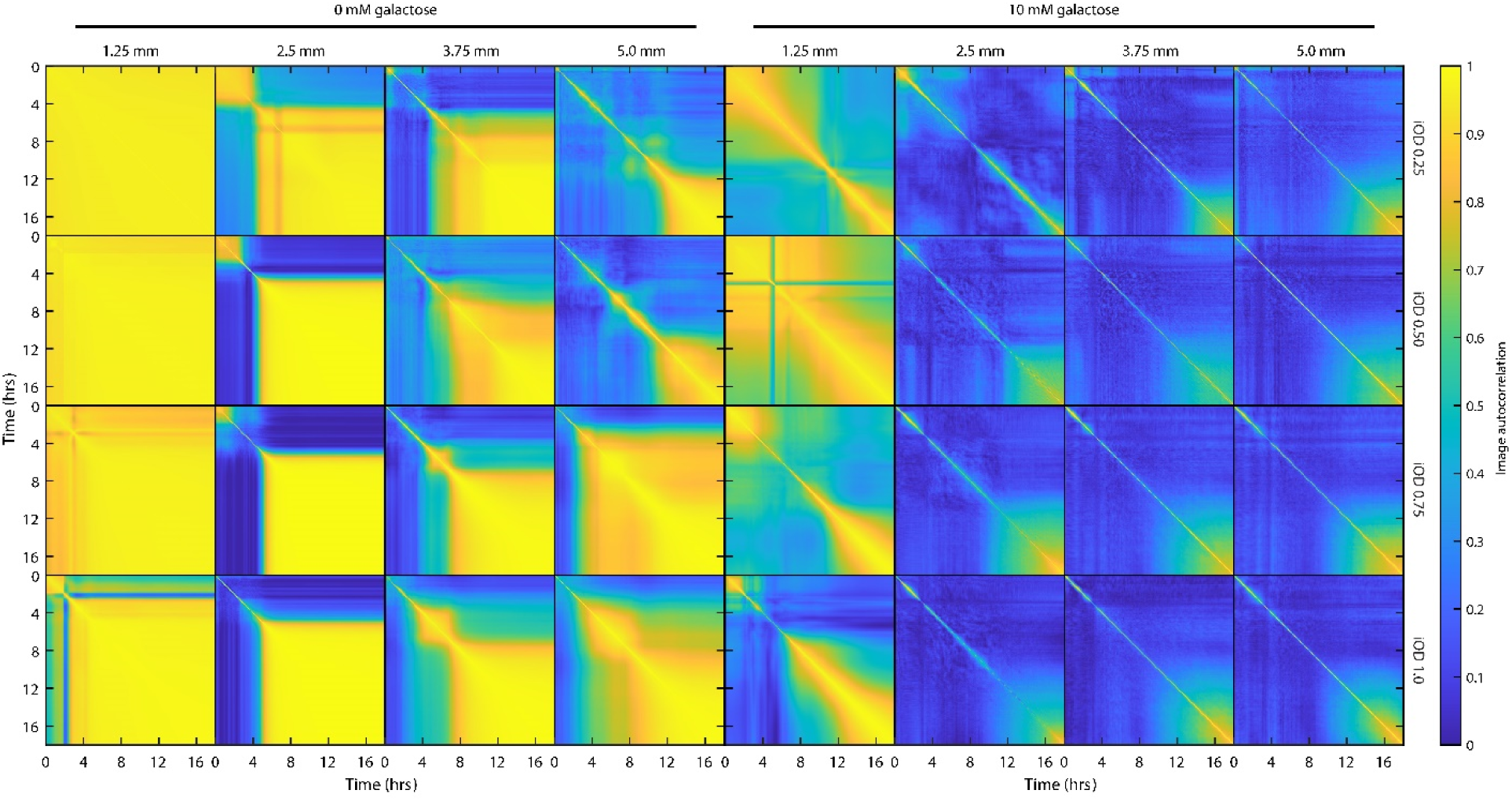
Cell-density correlation matrices as a function of fluid depth and initial OD in defined media with or without galactose. Each pane shows the cell-density correlation matrix for a particular fluid depth and initial OD, with either 0 mM or 10 mM galactose. Readily available carbon significantly increases the duration of bioconvection for depths greater than 1.25 mm (i.e., note far more blue on the right than the left); and for 1.25 mm, addition of galactose facilitates bioconvection, whereas it is absent under the same conditions without galactose (i.e., the large regions of yellow in the left-most column). All matrices are means over three individual experiments.

**SI Figure 7.**
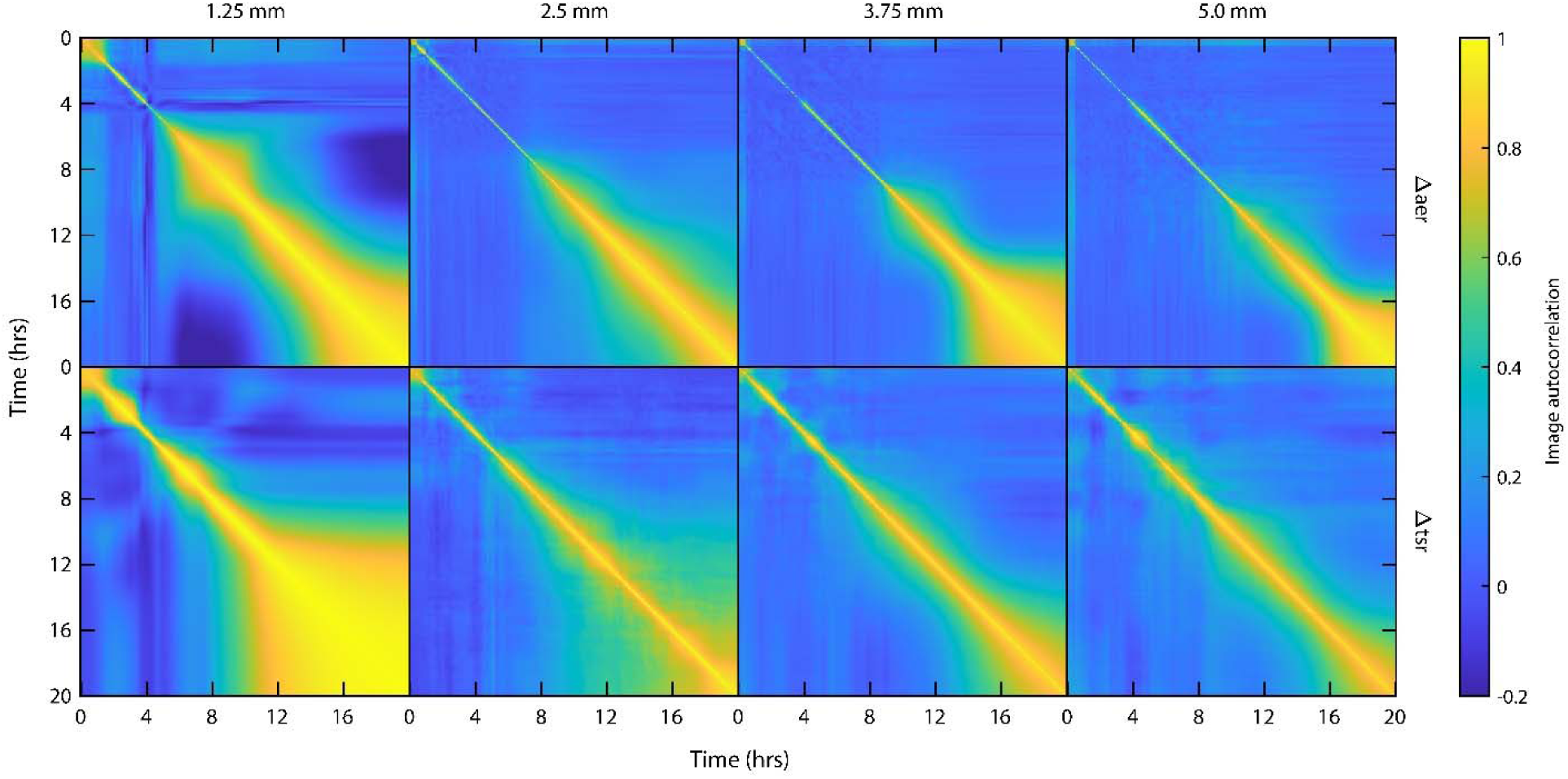
Cell-density correlation matrices for aerotaxis- and ‘energy-taxis’ -deficient mutants. Each pane shows the cell-density correlation matrix for a particular fluid depth for either aerotaxis-deficient mutants (top row) or energy-taxis-deficient mutants (bottom row). Whereas aerotaxis-deficient strains show similar patterns and timing trends to wild-type, energy-taxis-deficient strains display distinct patterns (see corresponding SI movies) and slower pattern evolution, at least in part due the emergence of slowly evolving sedimentation patterns. However, both the SI movies and the correlation matrices show comparably vigorous bioconvection in the bulk fluid above the sedimentation layer that lasts for >10 hours. All matrices are means over three individual experiments.

**SI Figure 8.**
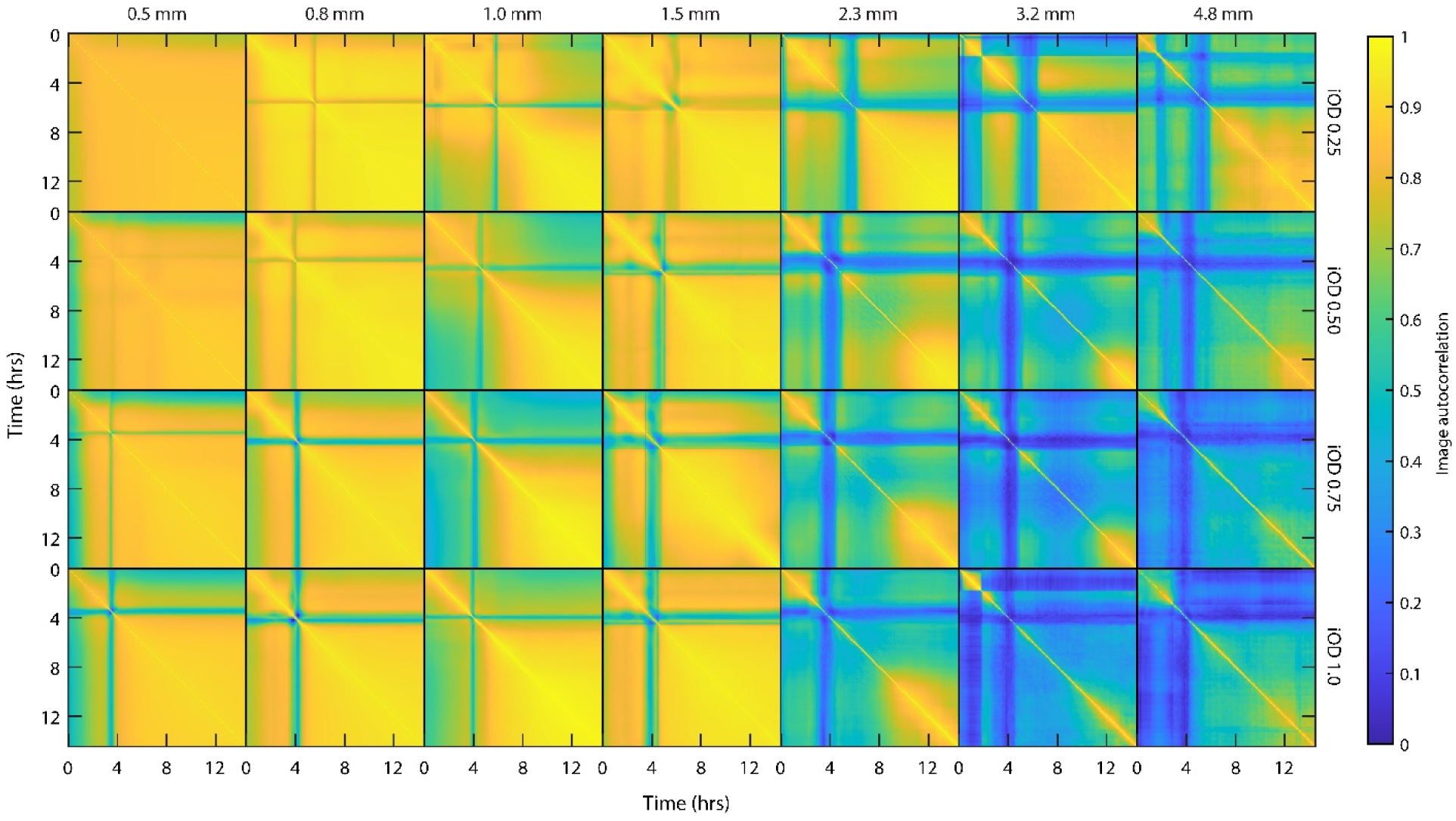
Cell-density correlation matrices across fluid depths in the absence of an air-liquid interface. Each pane shows the cell-density correlation matrix for a particular fluid depth and initial OD without an air-liquid interface. Similar to experiments with an interface, increasing fluid depth increased duration of bioconvection, and increasing initial OD decreased the onset time. However, whereas decreasing initial OD increased the duration of bioconvection in samples with an interface, here, decreasing initial OD tended to decrease the duration, with fluid depths below 2.3 mm displaying at most a relatively short-lived ‘pulse’ of bioconvection (∼1 hr). All matrices are means over three individual experiments.

## Notes

https://www.dropbox.com/s/j73iwaenr6r8j26/Figure_data_Shoup_Ursell_Bioconvection_2021_v1.zip?dl=1

### Competing Interest Statement

The authors have declared no competing interest.

